# Serotonin system development and adult function are regulated by GDNF

**DOI:** 10.1101/2024.12.11.627811

**Authors:** Daniel R. Garton, Isabel Runneberger, Laoise Casserly, Tabea Schorling, Ana R. Montaño-Rodríguez, Vilma Iivananen, Jaakko J. Kopra, Kärt Mätlik, Fredrik Fagerström-Billai, Anastasios Damdimopoulos, T. Petteri Piepponen, Jaan-Olle Andressoo

**Affiliations:** Department of Pharmacology, Faculty of Medicine & Helsinki Institute of Life Science, University of Helsinki, 00290 Helsinki, Finland; Division of Neurogeriatrics, Department of Neurobiology, Care Science and Society (NVS), Karolinska Institutet, 17177 Stockholm, Sweden; Bioinformatics and Expression Core Facility, Karolinska Institutet, Stockholm, Sweden; Department of Biosciences and Nutrition, Karolinska Institutet, Stockholm, Sweden; Division of Pharmacology and Pharmacotherapy, Faculty of Pharmacy, University of Helsinki, 00014, Helsinki, Finland

**Keywords:** Glial cell line-derived neurotrophic factor (GDNF), serotonin, serotonin system development, serotonin system function, neuropsychiatric disorders, mouse models

## Abstract

Cognitive functions, neuropsychiatric disorders and behaviors from feeding to mood relate to the serotonin (5-hydroxytryptamine; 5-HT) system. We report that expression of glial cell line-derived neurotrophic factor (GDNF), a known potent stimulator of the brain dopamine system, correlates with serotonergic markers in humans, and increased GDNF defines a subset of psychiatric patients with a characteristic 5-HT-related gene expression pattern. A similar ∼1.5- to 2-fold upregulation of endogenous GDNF expression in mice increases brain 5-HT levels and function, both developmentally and during adulthood, and modulates response to fluoxetine. Notably, increasing GDNF more than approximately 2-fold does not increase 5-HT further and instead produces an inverted U-shaped curve of 5-HT levels, suggesting why the GDNF/5-HT correlation has remained controversial. Collectively, our data indicate that GDNF levels fine-tune 5-HT system development and adult function while excess GDNF exclusively associates with neuropsychiatric illness, making it an important target for future research.

## 1 Introduction

Serotonin (5-hydroxytryptamine; 5-HT) has widespread roles in a variety of neuropsychiatric disorders and mental processes, including mood and affect,^1–3^ anxiety and stress,^4–6^ sleep,^5,7^ sexual activity, ^8^ and reward.^3,9^ As a common target of pharmacological interventions, 5-HT is implicated in human neuropsychiatric illness, including major depressive disorder (MDD),^10–12^ schizophrenia (SCZ),^13,14^ and bipolar disorder (BP).^15^ The 5-HT system innervates nearly every brain region^16^ including the striatum (STR) and two of the major dopaminergic neuron-containing structures in the brain: the substantia nigra (SN) and ventral tegmental area (VTA).^17,18^ To understand how 5-HT modulates neuropsychiatric illness and normal brain functioning, it is essential to determine what factors regulate 5-HT system development and adult function.^19^

The glial cell line-derived neurotrophic factor (GDNF) was first discovered as a potent neurotrophic factor for the growth and survival of dopamine (DA) neurons, with no significant effects observed on 5-HT using ectopic GDNF application.^20–23^ For this reason, ectopic delivery of GDNF has been tested in a bulk of preclinical trials in various Parkinson’s disease models where the nigrostriatal DA system is damaged, and in six clinical trials with additional clinical trials currently running.^24^ Historically, in terms of supporting the function of the DA system, however, clinical trials using ectopic GDNF have resulted in variable, inconclusive outcomes.^25^ The effect of GDNF on the brain 5-HT system on the other hand has remained controversial, with some studies claiming varying effects or increases to brain 5-HT,^26–32^ while others report little or no effect.^20–22,33–38^ An overview of these studies are presented in supplementary Table S1.

Moreover, how endogenous GDNF affects the brain 5-HT system development and/or adult function remains largely uninvestigated.^38^ However, recent single-cell sequencing studies revealed that serotonergic neurons of the dorsal raphe produce Ret mRNA,^39–42^ the receptor required for GDNF signaling in DA neurons,^43–45^ suggesting that these neurons can be responsive to GDNF.

Here, we first investigated the effect of GDNF on the brain 5-HT system using publicly available post-mortem gene expression data from SCZ, BP and MDD patients’ STR, prefrontal cortex (PFC) and hippocampus (HC).^46^ We found that high striatal GDNF expression levels exclusively associate with neuropsychiatric disorders and that GDNF levels correlate with the expression level of key 5-HT genes in the STR. We further define a uniform high GDNF/5-HT response group with a shared gene expression pattern in the STR.

To study causality, we then used mice where endogenous GDNF levels are elevated in the same range as in the GDNF/5-HT response group patients. We found that a relatively small 1.5- to 2-fold increase, similar to patients, in striatal GDNF level is sufficient to induce changes in 5-HT system development and adult function. Since drugs of abuse such as methamphetamine acutely increase nigrostriatal GDNF expression in the studied range in several species,^47–50^ we investigated an adult-onset increase in endogenous GDNF expression and observed induction of the 5-HT-specific transcription factor PET1,^51–54^ enhanced 5-HT related gene expression, and enhanced 5-HT levels. Similarly, in patients, GDNF levels correlated with expression levels of the human ortholog of PET1, FEV.^55,56^ Notably, the stimulation of 5-HT system function by GDNF is level-dependent, with an increase in GDNF greater than approximately 1.5- to 2-fold over wild-type activating a feedback loop by suppressing the expression of the rate-limiting enzyme in 5-HT synthesis tryptophan hydroxylase 2 (TPH2).^57,58^

## 2 Materials and Methods

### 2.1 Animals

Animal experiments were carried out in accordance with the European Union Directive 86/609/EEC. Additionally, the Regional State Administrative Agency of Southern Finland (license numbers ESAVI/11198/04.10.07/2014 and ESAVI/12046/04.10.07/2017) had approved all experiments. The number and suffering of animals was minimized to the greatest extent possible. At a relative humidity of 50-60 % and room temperature of 21±1 °C, the mice were group-housed in a specific pathogen free environment in individually ventilated cages with *ad libitum* access to food and water under a 12-h light-dark cycle (lights on at 6 a.m.). There were weekly bedding (aspen chips, Tapvei) and nest material (Tapvei) changes. Wooden blocks (Tapvei) were placed in cages for enrichment.

Previously generated GDNF knockout (GDNF cKO),^59^ GDNF constitutive hypermorph (GDNF hyper),^60^ and Cre-inducible GDNF conditional hypermorph (GDNF cHyper)^61,62^ mouse lines were used in the study. Nestin-Cre mice were B6.Cg-Tg(Nes-cre)1Kln/J mice obtained from the Jackson Laboratory (RRID:IMSR_JAX:003771). Wild-type littermates were used as controls in all experiments. Mice were maintained in a 129Ola/ICR/C57bl6 mixed genetic background. Mice were considered adult in the age range of 3-10 months. The researchers performing all behavioral tests, tissue collection and processing were blinded to the animal genotypes.

### 2.2 Home Cage Activity

Male mice were single-housed in InfraMot boxes for continuous daily activity monitoring. Mice were provided *ad libitum* access to food throughout the experiment and water for the first week. Water intake was monitored by the experimenter. Then, for the fluoxetine experiment, mice were administered 15mg/kg/day fluoxetine (Sigma). The concentration of fluoxetine in drinking water was adjusted to the estimated water intake. New fluoxetine solutions were prepared in every 2-3 days. Mice were kept on fluoxetine for 2 weeks, followed by a 1 week of rebaseline with water again *ad libitum*.

### 2.3 Dissections

Animals were deeply anaesthetized with CO_2_ and/or cervically dislocated and immediately decapitated. The skull was cut longitudinally open and the brain removed and placed into ice-cold phosphate-buffered saline (PBS) prior to tissue isolation. 2-mm thick coronal slices were obtained using a brain matrix (Stoetling), and brain structures of interest were then dissected from these slices and immediately snap frozen using dry ice. Sampled were stored at −80°C until use for HPLC or RNA extraction.

### 2.4 Histological analysis and immunohistochemistry

Mice were transcardially perfused for 2-3 minutes with warmed PBS at a flow rate of 10- 12 ml/min followed by warm 4% paraformaldehyde (PFA) (Thermo Fisher Scientific) for 5 minutes. Adult brains were post-fixed for up to 24 hours with 4% paraformaldehyde and cryosectioned after 2-3 days of incubation in 30% sucrose (Thermo Fisher Scientific) in PBS. Immunohistochemistry was performed on freely floating serially sectioned 25-μm coronal cryosections, stored at −20 °C in cryopreservant containing 30% ethylene glycol and 20% glycerol in 0.1M phosphate buffer, such that every 6^th^ section was stored together. The primary antibody used for all stains was an anti-5-HT antibody raised in rabbit (Sigma Aldrich, S5545), with an AlexaFluor 594 conjugated donkey anti-rabbit secondary antibody (Abcam, ab150064).

### 2.5 Stereology

Unbiased stereological cell counts were performed in a similar manner as described previously.^61^ Immunostained cryosections containing the dorsal raphe from 8-11 male mice per experimental group (approximately −4.04 to −5.2 mm AP from Bregma, according to the brain atlas from Paxinos & Franklin^63^) were sectioned in serial to a thickness of 25µm such that every 6^th^ section was taken for stains spanning the whole dorsal raphe. Cell bodies of 5-HT positive neurons in the dorsal raphe region were identified using a Zeiss microscope outfitted with MBPF stereology software. Stereological counts were done to ensure a Gunderson coefficient of error (m=1) less than 0.1 to estimate precision, and the counting grid was randomized by the software to cover 10 percent of the area of interest.

### 2.6 RNA extraction

Brain tissue was manually homogenized in Eppendorf tubes containing Trizol Reagent (Thermo Fisher Scientific) to extract total RNA from the samples according to manufacturer instructions, as described previously.^64^ Samples were kept on ice as much as possible. RNA quality was checked using a NanoDrop ND-1000 Spectrophotometer (Thermo Fisher).

### 2.7 Reverse transcription and quantitative PCR

Complementary DNA was generated from RNA as described previously.^64^ 200 to 500 ng of total RNA per sample was treated with RNase-free DNase I (Thermo Fisher Scientific). DNase I was inactivated with 5 mM EDTA at 65 °C, immediately followed by a reverse transcription reaction using random hexamer primers and RevertAid Reverse Transcriptase (Thermo Fisher Scientific) to yield complementary DNA (cDNA). cDNA was diluted 1:10 and stored at −20 °C until used for quantitative PCR (qPCR).

Quantitative PCR was performed as previously done^64^ with the BioRad C1000 Touch Thermal Cycler upgraded to CFX384 System (BioRad). Samples were supplied with SYBR Green I Master (Roche) and 250 pmol primers to make 10 μl total reaction solution volume pipetted into 384-well plates. Each reaction well included cDNA or a negative control consisting of either a no reverse transcription control or water. Both negative controls were always included on each plate, and all samples and controls were analyzed in duplicates. Mouse Rn18s, ActinB and Gapdh were used as reference genes. Primer sequences are given in the supplementary Table S2. Results for a biological repeat were discarded when the C_q_ value for one or more of the replicates was 40 or 0, or when the C_q_ difference between replicates was >1.

### 2.8 Genotyping

Extracta DNA Prep for PCR - tissue (Quanta Biosciences, USA) was used to isolate DNA from tissue samples taken from the tail or ear, with samples collected at mouse weaning and/or during dissections. AccuStart II GelTrack PCR SuperMix (Quanta Biosciences, USA) was used for DNA amplification and analyzed by electrophoresis using 2% agarose gels in TAE buffer (Elatus Media Kitchen, University of Helsinki). Primers used for genotyping the GDNF conditional knockout allele, the GDNF hypermorph allele were, and for the GDNF conditional hypermorph allele are given in the supplementary Table S3.

### 2.9 High-Performance Liquid Chromatography

High-performance liquid chromatography (HPLC) with electrochemical detection was used to identify and quantify 5-HT and its metabolite levels as described previously.^38,61^

### 2.10 Stereotaxic surgery

Intrastriatal AAV5-Cre viruses were produced and used as described previously.^61,64^ The mice were 3-5 months old at the time of the injections. Wild-type littermate controls were injected with the same dose of AAV5-Cre as the GDNF cHyper heterozygous and homozygous mice. Mice were anesthetized in 100% oxygen with isoflurane (3-4% for induction and 1.5-2% for maintenance; Oriola, Finland). The top of the head was shaved and Desinfektol P (Berner Pro, Finland) was used to disinfect the skin. Mice were placed on a stereotaxic surgery apparatus and the local anesthetic lidocaine (Yliopiston Apteekki, Finland) was injected into the skin above the skull. The skin in the aforementioned area was cut open with a scalpel and holes were drilled into the skull bilaterally. The coordinates used for the intrastriatal injections were A/P 0.7, M/L ±2.2, relative to Bregma. A 33G blunt NanoFil needle (World Precision Instruments, USA) was inserted at a 10- degree angle through the drilled holes into the brain until reaching D/V −3.0 mm relative to Bregma. 1.5 μl containing 1.7 × 10^11^ viral genome copies of AAV5-Cre diluted in Dulbecco’s PBS was injected into each site at a flow rate of 0.2 μl/min. The needle was kept in place for an additional 5 minutes prior to slow and careful retraction. The skin over the skull was then sutured shut. All animals were injected subcutaneously with 5 mg/kg carprofen (Yliopiston Apteekki, Finland) diluted in saline for post-operative analgesia. Sutures were examined the following day to ensure proper wound closure and healing.

### 2.11 Cyclic Voltammetry

6- to 8-month old male mice were decapitated and the brain was removed and placed immediately into oxygenated (95% O2, 5% CO2) ice-cold cutting buffer containing the following: 100mM glucose, 75mM NaCl, 2.5mM KCl, 26mM NaHCO3, 1.25mM NaH2PO4, 2mM MgCl2, and 0.7mM CaCl2. A model 7000 smz-2 vibratome (Campden Instruments) then cut slices in oxygen-bubbled (95% O2, 5% CO2) ice-cold cutting buffer. The brain was sliced coronally into 300-μm-thick slices containing the substantia nigra pars reticulata (SNpr) or dorsal raphe (DR). After slicing, the slices recovered in a holding chamber for 20 minutes at 34°C, and then for 1-4 h at room temperature in oxygen-bubbled (95% O2, 5% CO2) artificial cerebrospinal fluid (aCSF). The aCSF contained the following (in mM): 119 NaCl, 3 KCl, 26 MgSO4, 1 KH2PO4, 1.2 MgCl2, 2 CaCl2, and 10 glucose.

In the recording chamber at room temperature, the slices were perfused continuously with oxygen-bubbled aCSF. Fast-scan cyclic voltammetry recordings were performed using cylindrical 5 μm carbon fiber electrodes positioned at the SNpr or DR ∼50 μm below the top surface of the slice. Release of 5-HT was stimulated electrically using a stainless-steel bipolar electrode placed ∼100 μm from the recording electrode controlled by a Master-8 pulse generator (A.M.P.I.) which triggered and Iso-Flex stimulus isolator to produce square pulses 1 ms in duration. Slices were stimulated with 400µA of current. A stimulus magnitude of 400 µA was the minimum value that would reliably produce the maximum response. N-shaped voltage ramps from a holding potential of 0 mV up to +900 mV, down to −900 mV, and back to 0mV over 9 ms (scan rate of 400 mV/ms) were applied to the carbon fiber electrode at 100 ms intervals. An Axopatch 200B amplifier (Molecular Devices) was used to record the electrical current, which was filtered through a 5 kHz low-pass Bessel filter and digitized at 40 kHz (ITC-18 board; InstruTech). A computer routine in IGOR Pro (WaveMetrics) was used for N-shaped wave generation and data acquisition, and the recorded transients were characterized using the same software.^65,66^ Electrodes were calibrated using background-subtracted cyclic voltammograms obtained with 1 μM 5-HT solution diluted in PBS (5-HT; Sigma-Aldrich).

Serotonergic terminals were stimulated with a train of 30 electrical pulses at 30Hz over 1s at 2 min intervals, as described previously.^67,68^ In fluoxetine experiments, a concentration of 10µM fluoxetine (Sigma), sufficient to inhibit 5-HT but not dopamine reuptake,^69^ was added to the aCSF solution and perfused over the slices after a stable recording baseline of 3 peaks was achieved.

### 2.12 Statistical Analysis

Statistical analyses regarding the levels of GDNF in human patients, human gene expression correlations, and all mouse data except the RNA-sequencing were performed using GraphPad Prism version 10. Potential outliers were excluded if determined to be an outlier by Grubbs’ Test with an alpha of 0.05. Datasets were tested for normality and then significant differences were determined using t-test, one-way, or two-way ANOVA with or without repeated measures where appropriate. Statistical analyses pertaining to bioinformatics analyses can be found in the relevant sections below.

### 2.13 Data accession, pre-processing and differential gene expression analysis

RNAseq data from GDNF cHyper mice has previously been published by our group in Mätlik et al.^61^ and is accessible via GEO: GSE162974. Fastq files were aligned to mouse reference genome mm10 with STAR (RRID:SCR_004463) and annotated exons were counted by featureCounts (RRID:SCR_012919). Differential gene expression analysis was performed in R, using genes package edgeR (RRID:SCR_012802). Only genes with less than one count per million (CPM) in at least three of the samples were sifted out and for the remaining genes, TMM normalization, trended with dispersion estimation was used. The cut-off for significance was set to FDR = 0.1.

Microarray data from post-mortem STR, PFC and HC has been published by Lanz et al.^46^ and we gained access to the raw data via GEO: GSE143136. The samples were pre-processed using the package oligo (RRID:SCR_015729) in R followed by annotation with affycoretools.^70^ Differential gene expression analysis was performed with edgeR and linear modelling fitting with *eBayes* in limma^71^ using only genes with annotated entrez ids. Significance cut-off was set to FDR = 0.1 for the full disorder groups and to FDR = 0.05 for the GDNF/5-HT response group. GDNF expression in the post-mortem patient samples were plotted with ggplot2 (RRID:SCR_014601).

### 2.14 Serotonin genes in patient samples and GDNF cHyper mice

A list of 5-HT-related genes was created by combining the translation of 5-HT-related GO terms back to genes using the database org.Hs.egGO2ALLEGS with a list of 5-HT genes pulled from relevant 5-HT-related gene sets published in the Molecular Signatures Database^72–75^ and identified in animals from Mätlik et al.^61^ The chord diagram visualization of 5-HT genes in the GDNF/5-HT response patient group and GDNF cHyper mice was created with circlize (RRID:SCR_002141). For comparison between species, a list of entrez ids and gene symbol homologues between humans and mice was created using NCBI’s Ensembl annotations. Venn diagrams were visualized with VennDiagram (RRID:SCR_002414). The UMAP of expression of 5-HT genes in patients was created with uwot^76^ and supervised by disorder. Hierarchical clustering of patients using the 5-HT genes was performed with *hclust* and method ward.D2, and visualized via factoextra (RRID:SCR_016692). Heatmaps of 5-HT gene expression in patients were created with ComplexHeatmap (RRID:SCR_017270) after expression value standardization to mean equals 0 and standard deviation equals 1. Gene set analysis was performed with the functions *ids2indices* and *camera* in limma^71^ and *tricubeMovingAverage* was used when plotting.

## 3 Results

### 3.1 Striatal GDNF levels correlate with psychiatric disorder and expression of key serotonin system related genes

To investigate whether there is a correlation between GDNF expression, psychiatric disorder, and 5-HT system related gene expression in humans, we reanalyzed post-mortem human neuropsychiatric patient mRNA expression data from the striatum (STR), hippocampus (HC), and prefrontal cortex (PFC) from Lanz et al. (2019)^46^ and generated heatmaps depicting normalized 5-HT system related gene expression levels in the STR (Figure 1A) and in the PFC and HC (Figure S1). We found that, GDNF levels in the STR, one of the major expression sites for GDNF in the brain,^77,78^ are significantly increased patients with SCZ and BP (Figure 1D), but that the levels vary within each disorder group, and that in all three neuropsychiatric disorders there are patients with high GDNF expression above the control range (Figure 1A, D). In the HC and PFC, where GDNF is not expressed in cell bodies,^77–79^ GDNF levels are comparable to healthy controls (Figure S2). We observed that striatal GDNF mRNA levels positively correlated with both 5-HT biosynthesis rate-limiting enzyme tryptophan hydroxylase 2 (TPH2) and the 5-HT reuptake transporter (SERT) mRNA levels in human STR (Figure 1B), with specific significant correlations being found between GDNF and TPH2 in SCZ and MDD patient groups and between GDNF and SERT in the SCZ patient group (Figure S3).

**Figure 1.**
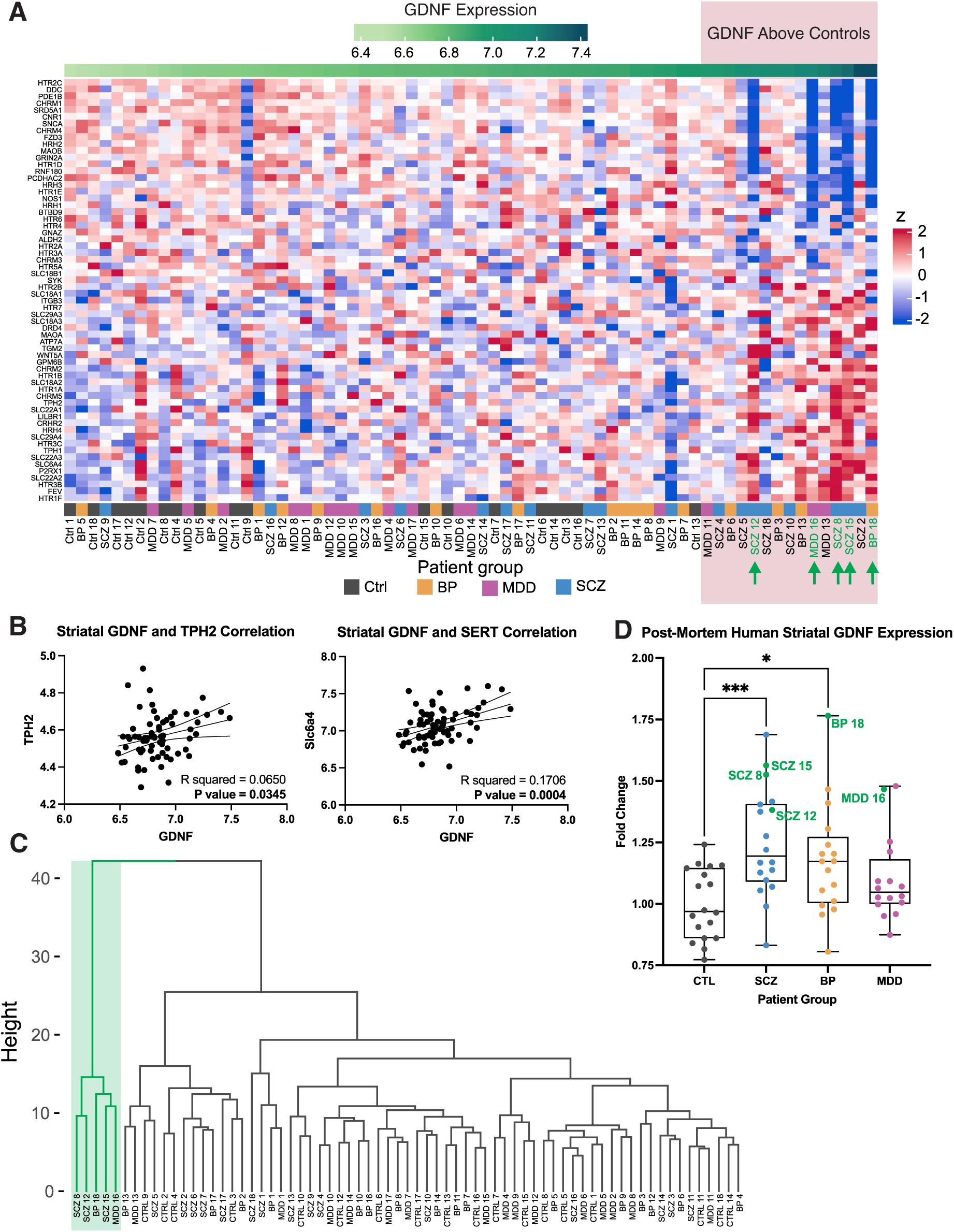
Increased levels of striatal GDNF associate with a GDNF/5-HT response group of neuropsychiatric patients. (A) 5-HT gene set expression heatmap from human postmortem striatal mRNA microarray data published in Lanz et al. 2019, with patient & control samples organized by ascending levels of GDNF expression. Red box denotes patient samples with GDNF level above control sample range. (B) In human striatal samples from Lanz et al., 2019, GDNF mRNA level linearly correlates significantly with both TPH2 and SERT mRNA expression. (C) Hierarchical clustering of the expression data depicted as a heatmap identifies subset of 5 human patients (3 SCZ, 1 BP, and 1MDD; green highlight) with similar striatal 5-HT gene expression pattern response. (D) GDNF mRNA levels is significantly increased in BP and SCZ compared to controls from Lanz et al., 2019 microarray data (one-way ANOVA p=0.0022; Holm-Šídák post-hoc test *** = p<0.001, * = p<0.05). Green labelled data points indicate 5 patients identified in patient subgroup from (C) with similar 5-HT gene set expression pattern, all five patients with notably high levels of GDNF above control range.

Heatmap analysis suggested that some patients with high GDNF levels display a distinct expression pattern of 5-HT genes (Figure 1A, green arrows). We employed a hierarchical clustering method to these data to investigate whether this unique expression pattern defines a patient subgroup and found a subgroup of five patients, deemed the “GDNF/5- HT response group”, across the three disorder categories (Figure 1C), all with high GDNF levels above controls (Figure 1D; marked with green). UMAP analysis confirmed that the “GDNF/5-HT response group” makes up its own distinct subgroup cluster (Figure S4).

### 3.2 Analysis of the role of the increased GDNF on serotonin system gene expression in mice

After establishing that high levels of GDNF are associated with neuropsychiatric illness, upregulation of TPH2 and SERT, and distinct changes in the 5-HT system, we next sought to determine causality: whether the relatively moderate, ∼1.5- to 2-fold increase in endogenous striatal GDNF levels observed in patients can itself be a cause of an altered 5-HT system. To this end, we next utilized mouse lines with elevated expression of endogenous GDNF protein and mRNA (Figure 2A): a GDNF Cre-inducible conditional hypermorphic (GDNF cHyper) line,^61,62^ and a GDNF constitutive hypermorphic (GDNF hyper) line.^60^ The GDNF cHyper mice display approximately a 2-fold and a 4-fold increase in endogenous GDNF expression in heterozygous and homozygous mice, respectively, starting from about embryonal day 12 (E12) onwards in the CNS after crossing to a Nestin-Cre line.^61,62^ In addition, the GDNF hyper mice have approximately a 2-fold increase in endogenous GDNF expression in heterozygous mice constitutively.^60^ As indicated in Figure 2A and described previously,^60–62^ in both mouse lines, the native 3’UTR of *Gdnf* is replaced, leading to increased expression at the post-transcriptional level, which warrants increased GDNF expression restricted to cells which natively transcribe *Gdnf* gene. Here, again, we demonstrate, respectively, a 2-fold and between a 3- to 4- fold increase in heterozygous and homozygous brain GDNF levels in GDNF cHyper mice already at P11 (Figure 2B). We also analyzed the effect of brain specific GDNF deletion on the 5-HT system using our previously generated GDNF conditional knock-out (KO) line (Figure 2A).^59^

**Figure 2.**
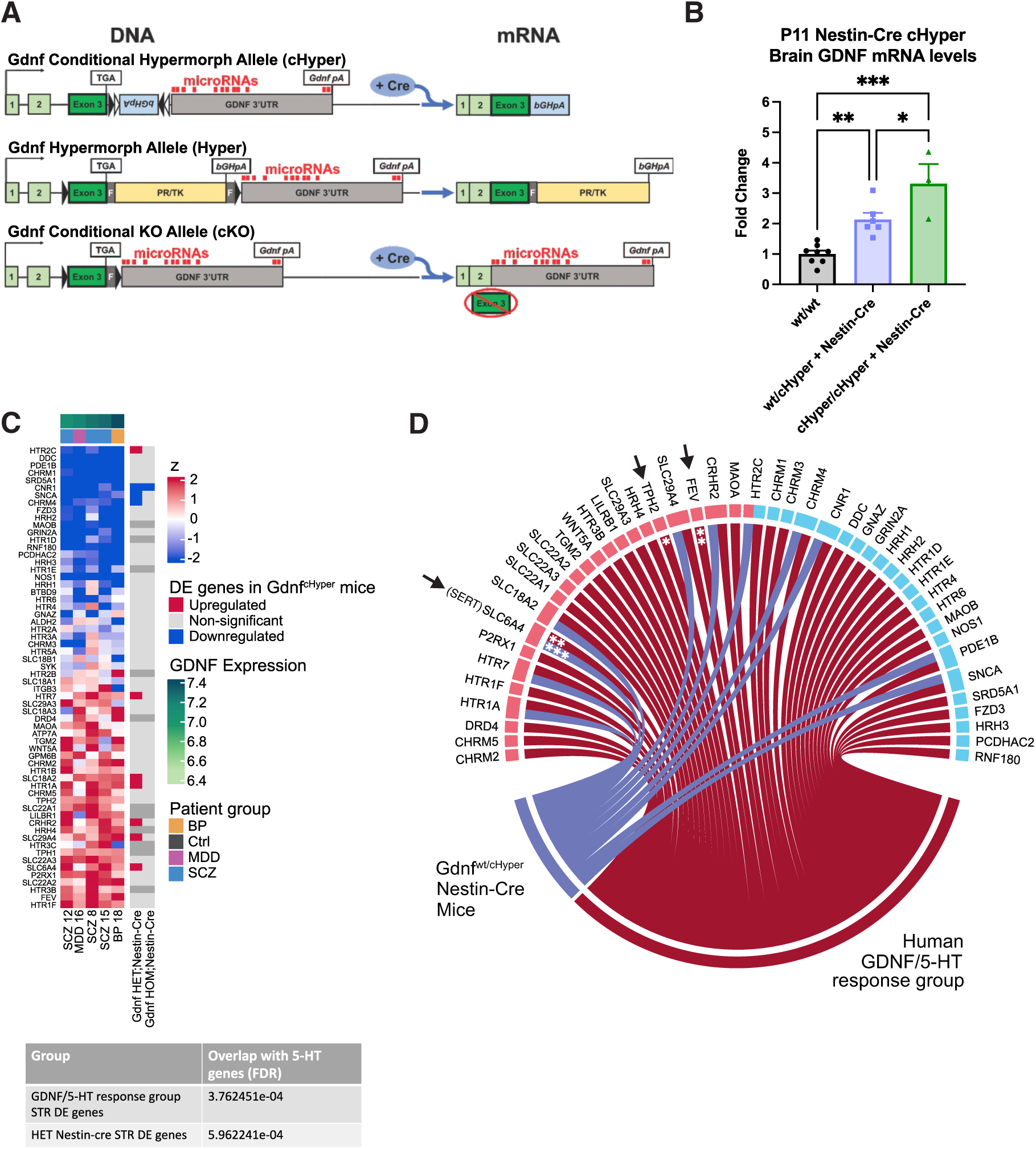
Human GDNF/5-HT response group gene expression pattern is similar to heterozygous Nestin-Cre GDNF conditional hypermorph animals. (A) Schematic representation of knock-in alleles used to target GDNF, with upside-down bGHpA floxed by LoxP sites which turns around the early polyA (pA) signal and shortens the 3’UTR in the presence of Cre-recombinase in conditional hypermorph animals (cHyper; top), inserted PR/TK cassette with early bGHpA signal, shortening 3’UTR, preventing microRNA binding and conferring mRNA stability in the hypermorphs (Hyper; middle), and floxed exon 3, corresponding to the region coding for the mature GDNF protein, for GDNF KO (cKO; bottom). Triangles represent LoxP sites. Grey boxes represent FRT sequences. (B) Brain GDNF mRNA level from Nestin-Cre cHyper mice increased significantly in a linear, stepwise manner according to number of cHyper alleles already at P11 (One-way ANOVA p<0.0001; Holm-Šídák post-hoc test *** = p<0.001, ** = p<0.01, * = p<0.05). (C) Striatal 5-HT genes expression heatmap from human GDNF/5-HT response group patients and from Nestin-Cre GDNF wt/cHyper heterozygotes, but not homozygotes, significantly overlap with 5-HT genes as determined by a one-sided hypergeometrical test (p-values reported in table below heatmap, see methods for statistical details). (D) Cord diagram depicting significantly upregulated (red) and downregulated (blue) genes in the Nestin-Cre GDNF cHyper heterozygous animals (wt/cHyper) and the GDNF/5-HT response group of patients with similar 5-HT gene response pattern. Arrows point to significant upregulation of FEV, TPH2 and SERT, white stars indicate significance relative to wild-type animals (t-test followed by FDR correction *** = p<0.001, ** = p<0.01, * = p<0.05).

Comparison of striatal gene expression in the GDNF cHyper x Nestin-Cre animals to the 5-HT gene set in GDNF/5-HT response patients revealed a significant overlap in the 5- HT gene set list (Figure 2C), including a significant increase in SERT expression (Figure 2D). Interestingly, we found that the overlap is significant only in GDNF cHyper x Nestin-Cre heterozygotes with approximately a 2-fold increase in GDNF level, rather than in GDNF cHyper x Nestin-Cre homozygotes where the GDNF increase is between 3- and 4-fold over wild-type (wt) levels (Figure 2B). The human GDNF/5-HT respsonse group also had TPH2 and FEV striatal mRNA levels significantly upregulated (Figure 2D). We observed that in humans, 71% of the 65 5-HT genes in the gene set are up- or downregulated, and in GDNF cHyper heterozygote mice, 17% (Figure S5).

### 3.3 An approximate 2-fold increase in developmental GDNF level is sufficient to increase adult serotonin levels and function

Both in humans and rodents, the site of the highest TPH2 activity and the most *Tph2* mRNA is expressed in the dorsal raphe nucleus.^80–83^ We observed that in GDNF cHyper x Nestin-Cre animals, Tph2 mRNA in the dorsal raphe is significantly increased (Figure 3A). High Performance Liquid Chromatography (HPLC) analysis of 5-HT tissue levels in GDNF cHyper x Nestin-Cre mice revealed increased 5-HT in the ventral striatum (vSTR) and SN (Figure 3B). Notably, again in both qPCR and HPLC results, we observed the strongest increase in 5-HT levels and in Tph2 expression in heterozygous animals, whereas in the homozygous animals both Tph2 levels as well as 5-HT levels were not further increased, but even downregulated relative to the heterozygous animals. These results suggest the existence of a feedback loop where too prominent of an increase in GDNF levels trigger Tph2 and 5-HT downregulation compared to heterozygous mice (Figure 3A, B). We denote this phenomenon an excess GDNF-driven inverted U-shaped curve effect on the 5-HT system (Figure 3A-B).

**Figure 3.**
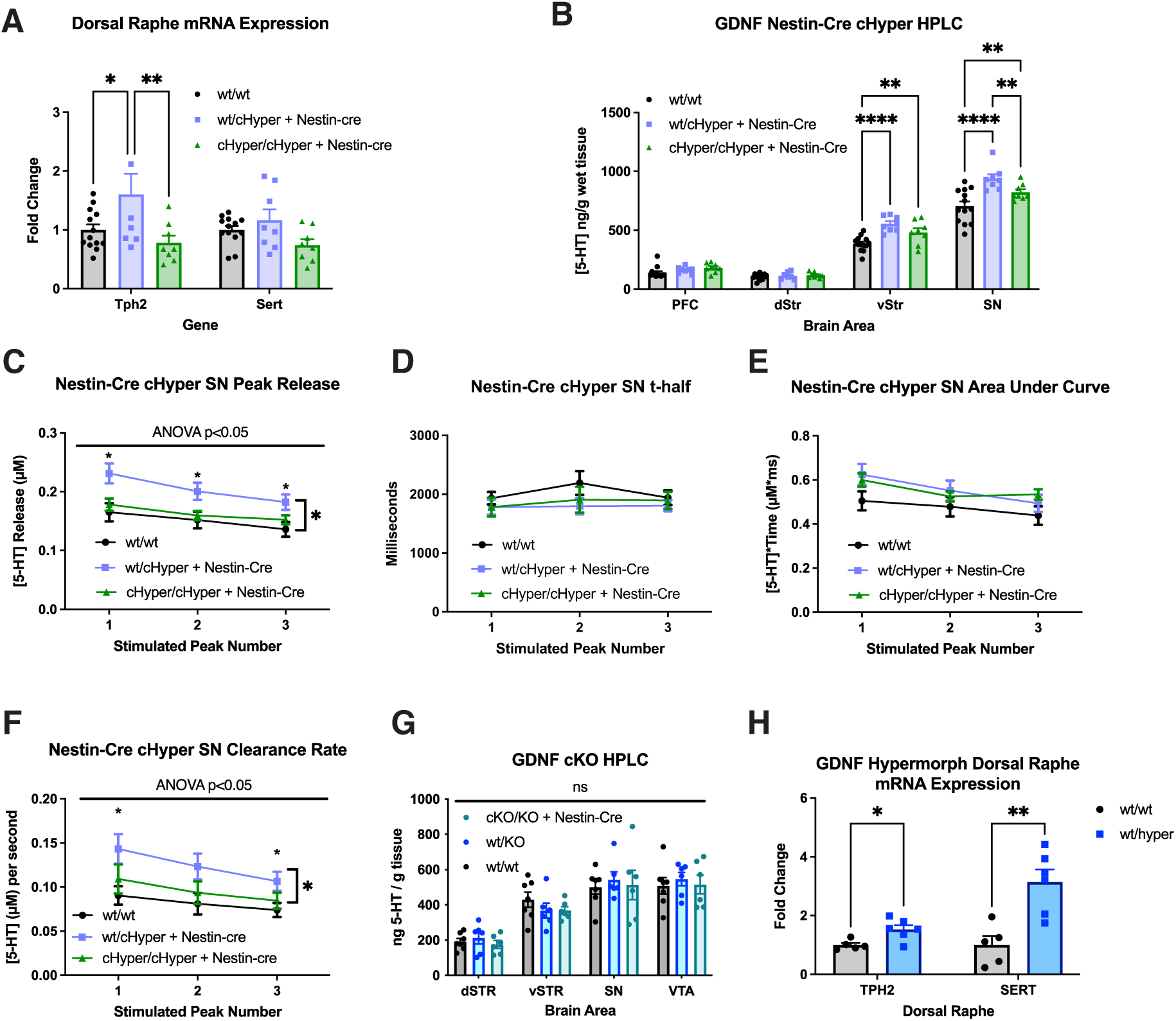
Only heterozygous mutant animals with an approximate 2-fold GDNF upregulation display increases in 5-HT levels, release, and reuptake. (A) Quantitative PCR of *Tph2* and *Sert* mRNA in the dorsal raphe of Nestin-Cre GDNF cHyper animals reveal a significant increase in Tph2 mRNA only in the heterozygotes, with a return to wild-type levels in homozygotes (two-way ANOVA indicates genotype as a significant source of variation; p = 0.0023, with Holm-Šídák post-hoc test indicating significant differences between groups; * = p<0.05; ** = p<0.01). (B) HPLC results in adult Nestin-Cre conditional hypermorphic animals show an increase in 5-HT concentration in heterozygous and homozygous animals in the vSTR and SN as compared to wild-type animals. In the SN, the homozygous animals have a decrease in 5-HT concentration as compared to heterozygous animals, indicating an inverted U-shaped curve effect. ANOVA shows significant interaction (**** = p<0.0001), with Holm-Šídák post-hoc test showing significant differences between genotypes in specific brain regions (** = p<0.01; **** = p<0.0001). N = 7-13 mice per group. (C-F) Fast-scan cyclic voltammetry from the SNpr of Nestin-Cre GDNF cHyper animals. (C) Electrically-evoked peak release of 5-HT was significantly increased in wt/cHyper heterozygotes as determined by a two-way repeated measures ANOVA with genotype as a significant source of variation, followed by a Holm-Šídák post-hoc test, indicating a main genotype effect between wild-type and heterozygotes (comparisons between all three genotypes; * = p<0.05) and simple effects at each electrically stimulated release (comparisons between wt and het; * = p<0.05). (D) The time it takes for electrically evoked release to reach half of its maximum (t-half) was unchanged between genotypes. (E) The area under the curve of the peak release event was unchanged between genotypes. (F) The clearance rate of 5-HT was also significantly increased in wt/cHyper animals (two-way repeated measures ANOVA revealed genotype as a significant source of variation; Holm-Šídák post-hoc test indicated a main genotype effect between wild-type and heterozygotes with comparisons between all three genotypes; * = p<0.05, and significant simple effects at electrically stimulated release peaks 1 and 3 with comparisons between wt and het; * = p<0.05). (G) HPLC of 5-HT levels in serotonergic neuron projection targets reveal no significant changes in 5-HT levels in adult GDNF knockout (cKO) animals. N=6-7 animals per genotype. (Two-way ANOVA p>0.05) (H) Increased levels of *Sert* and *Tph2* mRNA as measured by qPCR in the Dorsal Raphe of adult GDNF hypermorphs. Multiple student’s t-test corrected for multiple comparisons using the Holm-Šídák method reveals significant effect between genotypes (* = p<0.05, ** = p<0.01).

Next, we analyzed the function of the 5-HT system in GDNF cHyper x Nestin-Cre mice. Monoamine measurement using fast-scan cyclic voltammetry (FSCV) in the substantia nigra pars reticulata (SNpr) has previously been shown to predominantly comprise 5-HT over dopamine or norepinephrine.^67,84,85^ Using the same method as in John et al., (2006),^67^ we measured 5-HT release and reuptake in the SNpr. In line with both the qPCR and HPLC results, we found an increase in the peak release of 5-HT only in heterozygotes but not in homozygous animals (Figure 3C). The time it takes for the peak to reach half of its maximum (t-half) was unchanged between genotypes (Figure 3D), as was the area under the curve (AUC) of the total release-reuptake event, although there is a potential trend towards an increase in the mutant animals (Figure 3E). Because a greater peak release in combination with a similar t-half implies a disproportionally fast reuptake in the heterozygous animals, we next determined the clearance rate of 5-HT by dividing peak concentration by t-half, as done previously.^61^ We found, as above, that heterozygotes had a significantly faster 5-HT clearance rate compared to wild-type and homozygous animals (Figure 3F).

Mice lacking GDNF (GDNF knock-out, KO) die at birth due to a lack of kidneys,^86^ precluding postnatal analysis. Brain-specific GDNF deletion using Nestin-Cre or AAV-Cre changes dopamine system function,^64,87^ but does not affect dopamine neuron numbers.^59^ To study how GDNF deletion affects the 5-HT system in the adult rodent brain, we crossed Nestin-Cre animals to GDNF-conditional KO animals. We found that 5-HT levels in both systemic KO heterozygous animals and in adult mice with full GDNF deletion in the CNS are not changed (Figure 3G).

### 3.4 Constitutive ∼2-fold upregulation of endogenous GDNF increases 5-HT neuron number in caudal principal dorsal raphe and increases 5-HT system function

Next, we analyzed heterozygous GDNF hypermorphic mice (GDNF hyper), where approximately a 2-fold increase in endogenous GDNF expression via 3’UTR replacement is constitutive.^60^ Homozygous GDNF hyper animals die within 1-2 weeks after birth due to severely underdeveloped kidneys and a malformed urogenital system^60,88,89^; therefore, only heterozygous mice were analyzed. We observed that GDNF hyper animals display significantly increased levels of Tph2 and Sert mRNA in the dorsal raphe (Figure 3H), similar to what was observed in the STR of GDNF cHyper x Nestin-Cre heterozygous animals and in GDNF/5-HT response group patients.

To analyze the effect of increased GDNF levels on 5-HT system development, we performed stereological analysis of the 5-HT+ cells in the dorsal raphe of adult animals (Figure 4A). We observed a trend towards an increase in the numbers of 5-HT+ neurons in the principal dorsal raphe, as defined in Ren et al.^41^ (Figure 4B). However, analysis of rostral and the caudal portions of the principal part of dorsal raphe separately revealed that while the rostral part had no significant increase in the number of the 5-HT+ neurons, the number of 5-HT+ neurons in the caudal section of the principal dorsal raphe was increased (Figure 4 A,B).

**Figure 4.**
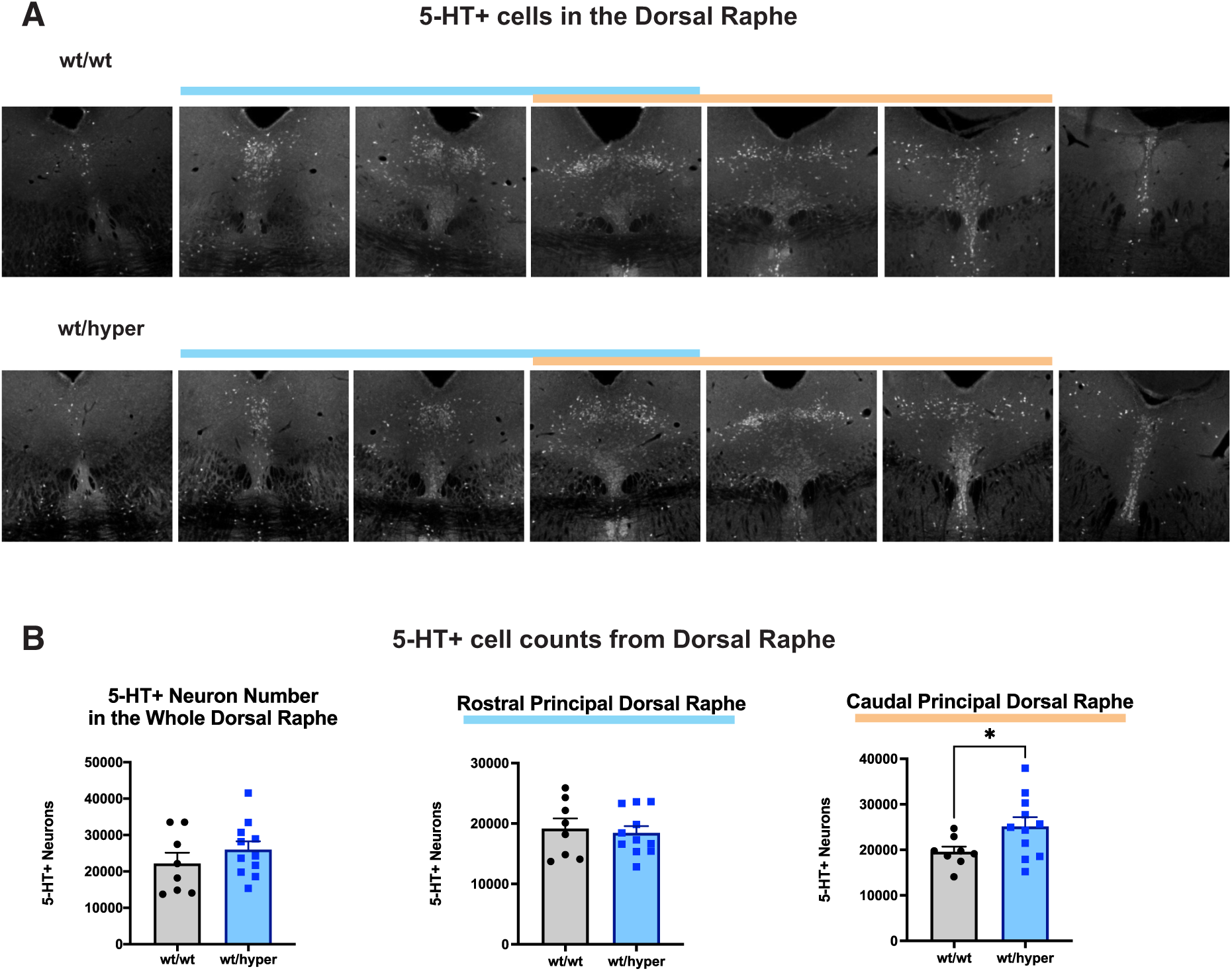
A developmental increase in GDNF in hypermorphic animals induces increase in 5-HT+ cell numbers in subsection of dorsal raphe (A) Representative images depicting 5-HT positive neurons stained with an antibody against 5-HT in the dorsal raphe of wild-type and GDNF hypermorphic animals. Blue line represents “rostral” portion of principal dorsal raphe, and orange line depicts “caudal” (B) Unbiased stereological cell counts of total 5-HT positive neurons in the total dorsal raphe (left), the rostral portion (middle), and the caudal portion (right). There was a significant increase in the number of 5-HT+ neurons in the caudal portion by unpaired t-test (* = p<0.05).

Consistent with the above cell count results, and in line with the GDNF cHyper heterozygous animals, in GDNF hyper mice, higher tissue levels of 5-HT were observed in the mutant animals across all midbrain and basal ganglia brain regions measured via HPLC (Figure 5A). To assess functionality, we next determined 5-HT release and reuptake using FSCV. In the SNpr, the evoked peak release of 5-HT and the AUC were significantly increased in the GDNF hyper animals (Figure 5B, D) while the t-half was unchanged (Figure 5C).

**Figure 5.**
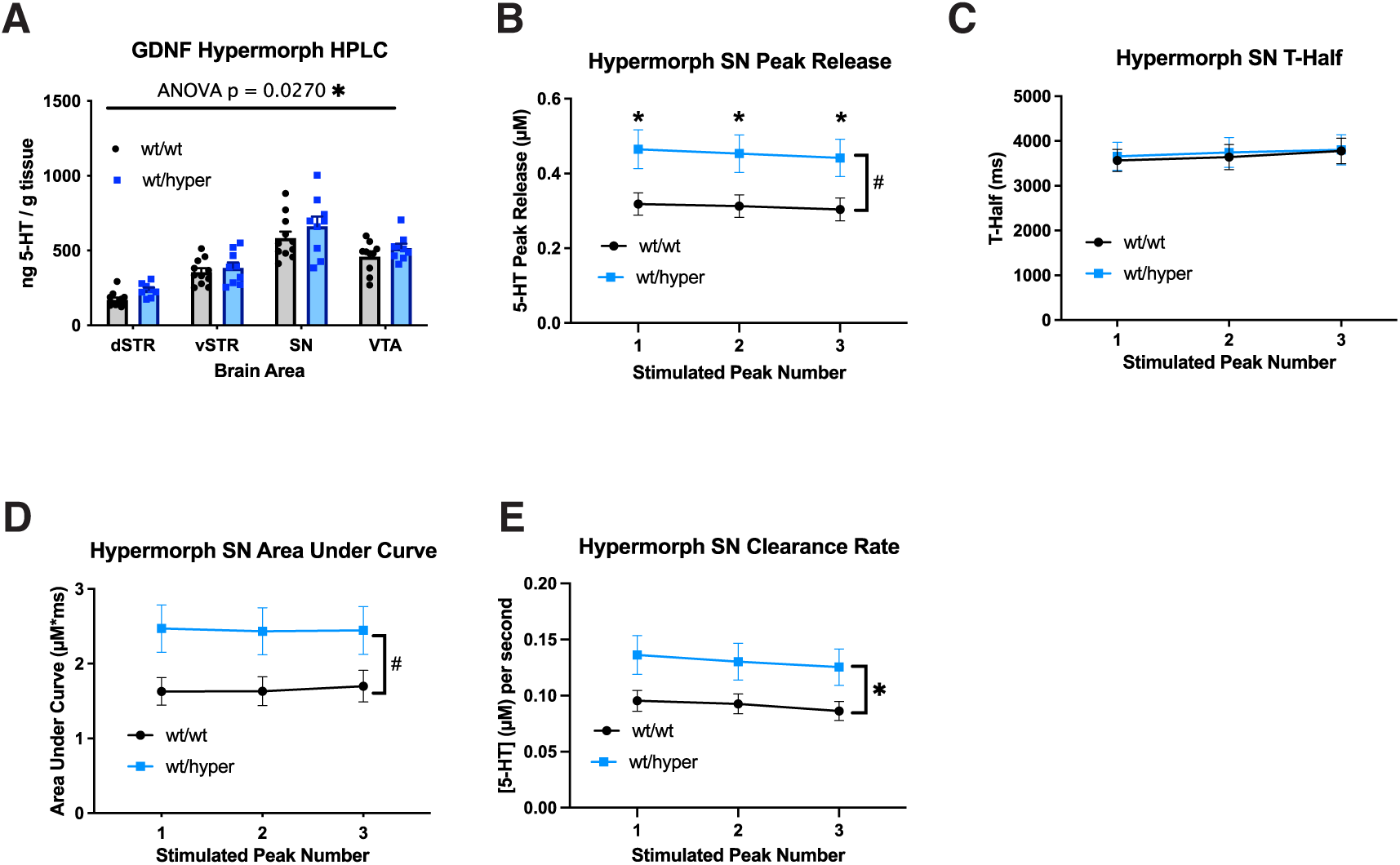
Heterozygous GDNF hypermorphic animals have increases in 5-HT levels, release, and reuptake. (A) HPLC results in adult animals show an increase in 5-HT concentration across all brain regions analyzed in GDNF hypermorphs. ANOVA shows significant effect of genotype (* = p<0.05). N = 8-11 mice per group. (B-E) Release and reuptake of 5-HT in the SNpr (B) Peak release in the SN is increased in adult hypermorphic animals. 2-way repeated measures ANOVA shows a significant effect of genotype (# = p < 0.05), and Holm-Šídák post-hoc test shows significant difference at each peak (* = p<0.05). N=18 nigral slices per genotype from n=6 mice. (C) No difference in observed t-half. (D) The area under the curve was significantly increased in the adult hypermorphic animals (2-way repeated measures ANOVA shows significant effect of genotype # = p < 0.05). (E) 5-HT clearance was significantly increased in the adult hypermorphs. 2-way repeated measures ANOVA shows a significant effect of genotype (* = p<0.05).

Like in GDNF-cHyper x Nestin-Cre heterozygous mice, there is a significantly increased clearance rate in the GDNF hyper animals in the SNpr (Figure 5E). FCSV analysis in the dorsal raphe revealed no significant differences between the genotypes in peak release, t-half, AUC, or clearance rate (Figure S6).

### 3.5 Increased GDNF increases response to fluoxetine

We found that GDNF hyper animals display an increase in serotonergic neuron number in the caudal principle dorsal raphe, in brain 5-HT levels, and in 5-HT release and reuptake in the SN. Thus, next we looked at how this affects their response to fluoxetine, a selective 5-HT reuptake inhibitor often prescribed to treat MDD and used as an auxiliary drug to treat other neuropsychiatric spectrum disorders.^90^ In *ex vivo* slices using FSCV, data was analyzed relative to the pre-fluoxetine stimulation baseline (white background) (Figure 6A-B). We found that 10µM fluoxetine when applied (purple background) increased t-half in both wt and mutant animals (Figure 6A), as previously reported,^67,69^ thus confirming that the FSCV measurements were indeed of 5-HT. Notably, the peak release in the GDNF hyper mutants increased significantly faster in response to 10µM fluoxetine (Figure 6B), rising above their normal baseline earlier than in the wt animals. However, at later stimulation time points, this apparent early increase in the GDNF hyper animals becomes comparable to the wt animals.

**Figure 6.**
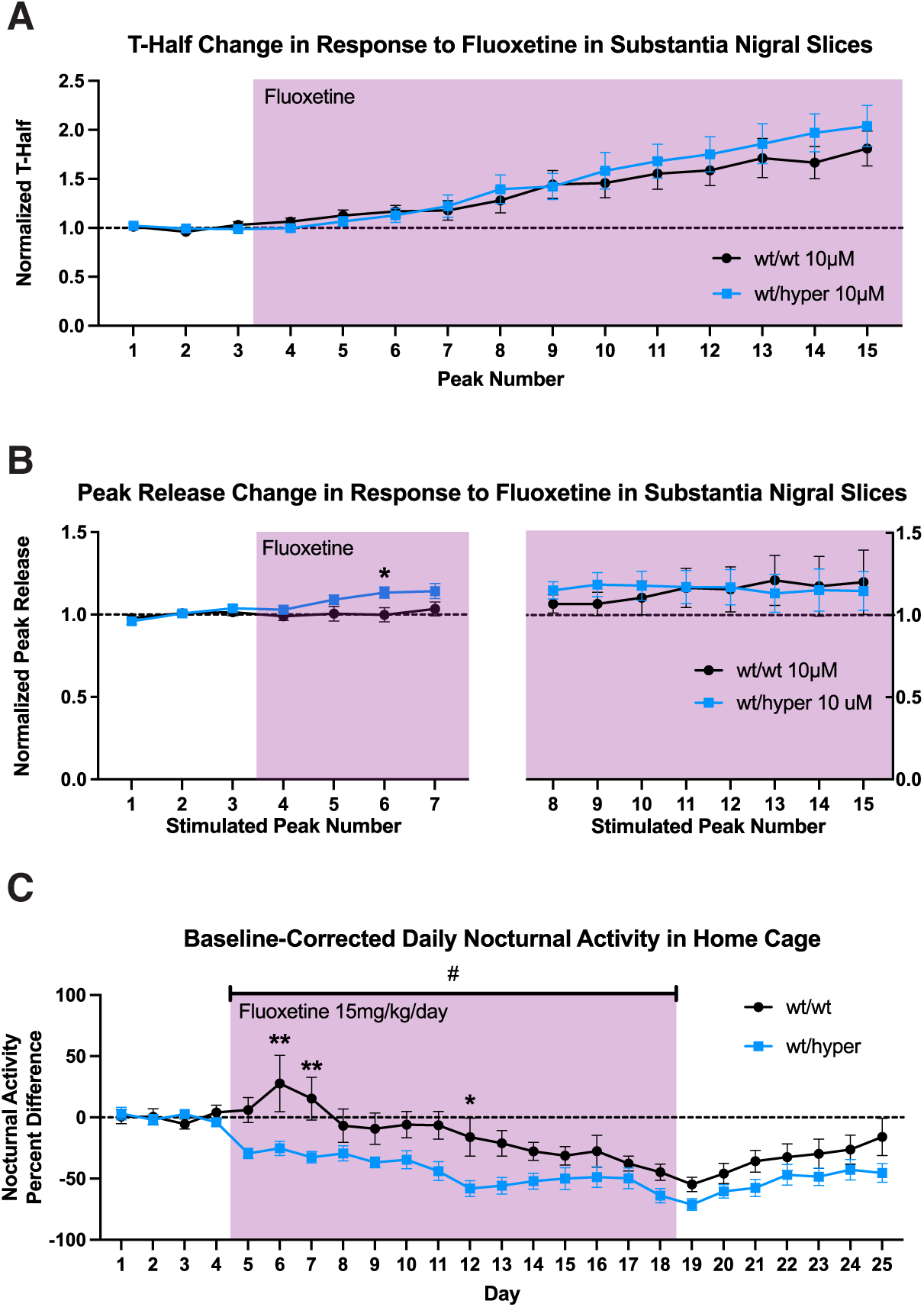
Heterozygous GDNF hypermorphic animals have increased response to fluoxetine. (A) SN slices from GDNF Hypermorphic animals respond to the SSRI fluoxetine showing a significant increase in t-half after fluoxetine is applied (purple shading), as expected from a competitive inhibitor, in both wild-type and hypermorphic animals in the area where recordings were done. Stimulation peaks are produced every 2 minutes, and the t-half is taken relative to the first three peak no-fluoxetine condition (average of the first 3 peak baseline is indicated with the dotted line). (B) Peak release is increased early after fluoxetine is applied (purple shading) in the Nigral slices from GDNF hypermorphic animals, however at later stimulation time points, the wild-type and hypermorph peak release are equally elevated above the first 3 peak no-fluoxetine condition (average of the first 3 peak baseline indicated by the dotted line). Two-way repeated measures ANOVA shows a significant interaction at early time points (p=0.019), with Holm-Šídák post-hoc test showing a significant difference at peak 6 (* = p<0.05), but not at later time points. (C) Baseline-corrected daily activity. Hypermorphic animals display an earlier, and increased activity reduction in response to fluoxetine. There is an overall ANOVA between genotype effect during fluoxetine treatment (# = p<0.05), as well as specific days significantly different as indicated by Holm-Šídák post-hoc test (* = p<0.05; ** = p<0.01).

Next, we analyzed whether GDNF hyper animals had an altered response to fluoxetine at the behavioral level by tracking their home-cage activity. Previous reports have demonstrated that chronic administration of fluoxetine reduces home-cage activity in mice.^91^ We similarly observed overall nocturnal activity of our mice is reduced by chronic administration of fluoxetine in both wild-type and mutant animals (Figure 6C, Figure S7). The GDNF hyper animals have a nonsignificant trend towards increased nocturnal activity prior to fluoxetine administration (Figure S7B). Thus, in order to measure the effect of fluoxetine on daily nocturnal activity, activity change was taken relative to each animals’ individual baseline (Figure 6C). Upon baseline-correction, the GDNF hyper animals display enhanced reduction in basal activity and a faster reaction to the fluoxetine, revealing an altered response to fluoxetine (Figure 6C).

### 3.6 Striatal GDNF increase in adulthood induces changes to the 5-HT system with a GDNF dose-dependent inverted U-shaped curve effect

GDNF is highly expressed in the STR,^79^ a known projection target of serotonergic neurons of the dorsal raphe, which also project to the SN via axonal collateral projections.^42,92^ Nigrostriatal GDNF levels are known to be induced, for example, by drugs of abuse including methamphetamine.^47–50^ To analyze the effect of an adult-onset increase in striatal GDNF we injected AAV-Cre bilaterally into the striata of adult GDNF cHyper animals (Figure 7A). Three months after bilateral intrastriatal AAV-Cre injections we observed on average about a 1.5-fold increase in Gdnf mRNA expression in heterozygous and about 2-fold increase in homozygous animals’ striata (Figure 7B). We again observed an increase in 5-HT levels as measured by HPLC in the vSTR and the SN (Figure 7C). Similarly, we again observed an inverted U-shaped curve of 5-HT levels, where homozygotes did not have a further increase in 5-HT levels compared to the heterozygous animals (Figure 7C). In line with the increase in 5-HT levels, there was a congruent significant increase in *Tph2* mRNA in the dorsal raphe of heterozygotes (Figure 7D), while homozygotes displayed no significant increase (Figure 7D), reflective of the inverted U-shaped curve of 5-HT levels (Figure 7C).

**Figure 7.**
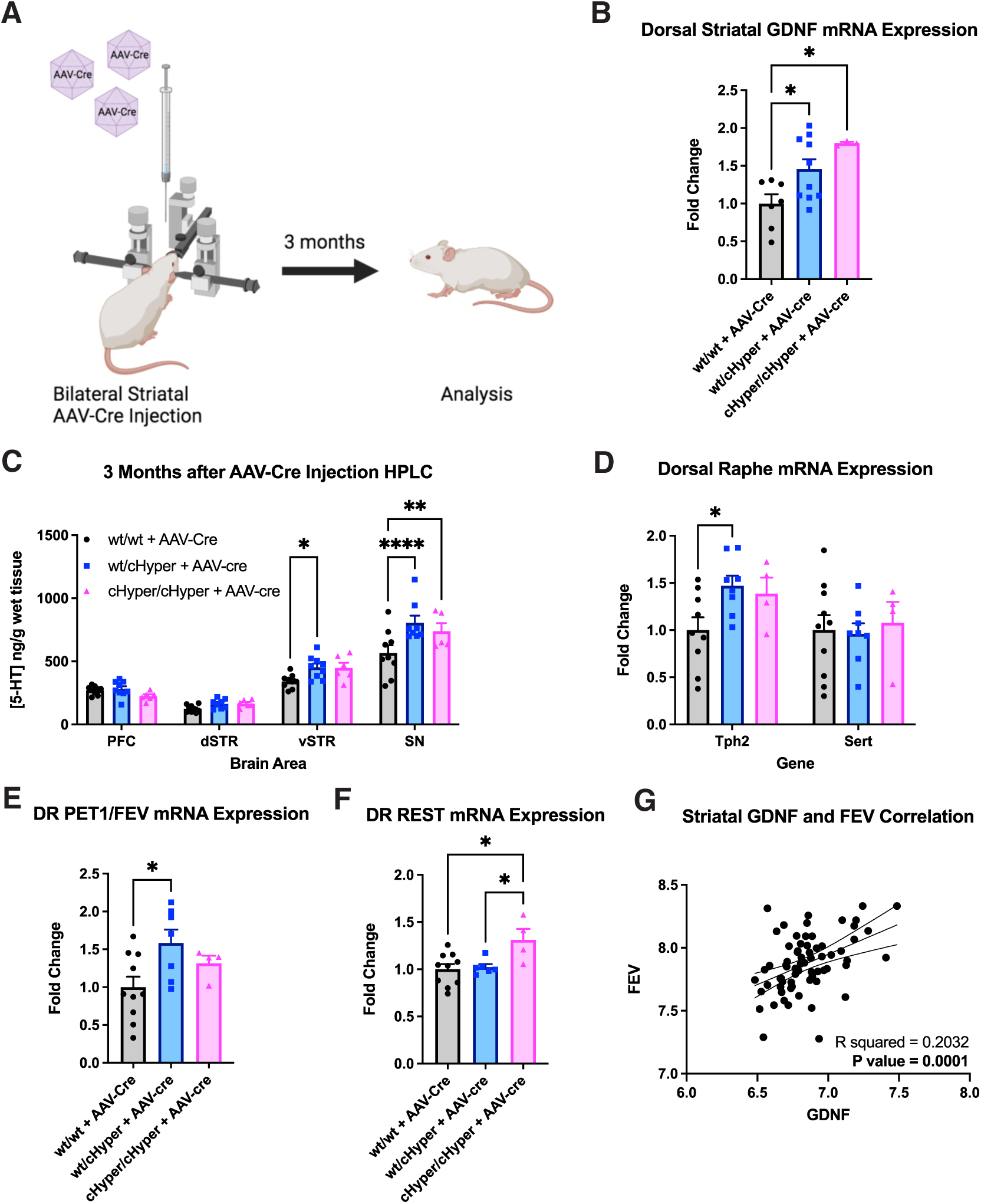
1.5-fold adult-onset striatal GDNF upregulation induces 5-HT system increase. (A) Experimental schematic depicting bilateral injection of AAV-Cre into the striata of adult mice. (B) Striatal GDNF mRNA level 3 months after injection shows significant, step-wise increase in GDNF levels (ANOVA between genotype p<0.05, Holm-Šídák post-hoc test * = p<0.05). (C) HPLC results from AAV-Cre injected animals of serotonergic neuron projection targets show increased 5-HT levels in the vSTR and SN of heterozygous animals, and in the SN of homozygous animals as compared to wild-type (ANOVA interaction p<0.01; Holm-Šídák post-hoc test reveals differences within specific brain regions). (D) *Tph2* and *Sert* mRNA levels in the dorsal raphe of AAV-Cre injected animals reveal a significant increase in *Tph2* in heterozygous animals, and a nonsignificant trend towards increase in homozygous animals (ANOVA interaction p<0.05; Holm-Šídák post-hoc test * = p<0.05). (E) Dorsal raphe Pet1 (orthologous to FEV in humans) mRNA expression is induced in heterozygous wt/cHyper animals by GDNF upregulation in the STR following AAV-Cre injection (ordinary one-way ANOVA p<0.05; Holm-Šídák post-hoc test * = p<0.05) (F). *Rest* mRNA in the dorsal raphe of striatally injected AAV-Cre animals is significantly increased in homozygotes (ANOVA p<0.05; Holm-Šídák post-hoc test * = p<0.05). (G) GDNF and FEV mRNA levels positively correlate significantly in human striatal sample data taken from Lanz et al., 2019 ^46^ (Pearson correlation r^2^ = 0.2032, p=0.0001).

Next, we analyzed the brain 5-HT system regulating transcription factor (TF) PET1, the TF well known to be required for the 5-HT system induction during development and maintenance during adulthood.^51–54^ PET1 induces Tph2 expression via direct binding to the *Tph2* promoter.^53,93,94^ We found that AAV-Cre mediated upregulation of striatal GDNF induces PET1 expression in the dorsal raphe in heterozygous but not in homozygous animals (Figure 7E).

Next, we analyzed whether the transcriptional repressor, REST, a known repressor of both *Pet1*^95^ and *Tph2*^96–98^ expression is changed by excess GDNF. We found that REST was upregulated only in the homozygous animals (Figure 7F), suggesting a mechanism for how excess GDNF triggers the inverted U-shaped curve in the brain 5-HT system in GDNF cHyper homozygous animals.

Next, we analyzed if and how an increase in striatal GDNF levels in humans, similar to the GDNF level increase in AAV-Cre GDNF cHyper heterozygous animals, correlates with FEV, the human orthologue to mouse PET1,^55,56,99^ expression. We found that GDNF and FEV mRNA levels were linearly positively correlated in the STR of humans (Figure 7G) with similar correlation observed in the PFC and HC (Figure S8).

## 4 Discussion

The 5-HT system is involved in numerous functions ranging from creative thinking and experiential processing^100^ to emotional regulation and mood (Reviewed in ^101^). For example, 5-HT release in the SN of humans has been recently directly shown using FSCV to track reward value,^102^ and high 5-HT levels modulate social behavior.^103,104^ Similarly, alterations in brain 5-HT levels play a role in neuropsychiatric illness including schizophrenia, bipolar disorder, and depression, which in some patients are comorbid and may reflect pathological continua with potentially similar underlying biological mechanisms.^105–109^ Thus, understanding what mechanisms regulate 5-HT system development and adult function has broad implications.

Here, using innovative conditional Hypermorph (cHyper) and constitutive Hypermorph (hyper) alleles, which result in an increase in expression limited only to cells which naturally transcribe Gdnf thus excluding ectopic expression driven artefacts (summarized in ^60,61^), we find that during development, GDNF levels regulate the number of 5-HT neurons in dorsal raphe (DR) nucleus, more specifically in the caudal part of the principal dorsal raphe defined in Ren et al. (2019).^41^ We further find that increased GDNF levels increase tissue 5-HT function by enhancing its levels in the brain most likely by inducing expression of Tph2, the rate limiting enzyme in 5-HT production in the brain,^57,58^ and by enhancing 5-HT release and reuptake at least in the SN, a brain region accessible for real-time 5-HT release and reuptake measurements.^67^

Thus, one mechanism which may explain individual differences in 5-HT-related functions may relate to variation in GDNF levels during development. Individual variance in GDNF levels may at least in part relate to the relatively long (2983bp in humans, 2500 bp in mice) and microRNA binding-site rich 3’UTR of GDNF, where fluctuations in individual miR levels can affect GDNF levels.^60,110^

Furthermore, here we find that an increase in endogenous GDNF level by as little as nearly 1,5-fold in adult STR, the main site of GDNF expression in the rodent brain,^77,78^ results in increased tissue 5-HT in the vSTR and SN. Notably GDNF is retrogradely transported to the midbrain from the STR by GDNF receptor RET expressing dopamine neurons and may be released in the midbrain where it may stimulate 5-HT axons projecting into the SN.^111,112^ Recent single-cell RNA-seq studies of the mouse brain have shown that at least a subcluster of serotonergic neurons in the dorsal raphe of mice express RET and project to structures of the basal ganglia, including into the STR and the SN.^40–42^ Thus, similar to DA neurons,^43–45^ GDNF is very likely to directly signal onto 5- HT neurons via RET. In parallel, 5-HT neurons which innervate the STR and receive the increased GDNF signal could upregulate 5-HT in the SN through 5-HT axon collaterals.^42,92,113^

Upon adult upregulation, GDNF induced the expression of transcription factor PET1 in the dorsal raphe. This is important since PET1 expression is restricted to 5-HT neurons,^51,114^ and it is required for 5-HT neuron phenotype generation and maintenance during adulthood.^52,53,115,116^ PET1 directly binds to the *Tph2* and *Sert* promoter regions to stimulate transcription,^53,93^ maintains chromatin accessibility to *Tph2* and *Sert* loci,^94^ and maintains serotonergic neuron connectivity and function in adulthood.^53,54^ It is thus conceivable that the observed increase in 5-HT function is mediated by a GDNF-induced increase in PET1 expression.

Importantly, we found that GDNF has a relatively narrow, concentration dependent, inverted U-shaped curve effect on 5-HT metabolism related gene expression and on 5- HT levels both during development and adulthood. This finding likely explains why previous reports on ectopic GDNF effects on the 5-HT system (summarized in Table S1) are contradictory with some reports claiming no effect^20–22,33–38^ while others report varying effects and/or stimulation.^26–32^ We find that at higher doses, an over 1.5-fold adult-onset striatal increase in GDNF induces REST, a known negative regulator of both FEV/PET1^95^ and TPH2 expression,^96–98^ suggesting a potential mechanism for how the observed inverted U-shaped curve manifests. It is, however, important to note that while brain 5-HT in homozygous animals returns towards wild-type levels, striatal dopamine, on the other hand, remains increased.^61^ Homozygous animals display a range of schizophrenia-like neurophysiological, neurochemical and behavioral deficits including apathy-like behavior, altered pre-pulse inhibition, and polydipsia along with increased striatal and reduced PFC dopamine.^61^ Thus, the inverted U-shape curve manifests on brain 5-HT, not on brain DA levels.

In line with the data from mice where only one parameter—GDNF level—is changed, in the human patients, we observe a correlation between mRNA levels of GDNF and FEV, the human orthologue of PET1,^99^ in all brain areas studied. FEV is also specifically expressed only in serotonergic neurons of the midline brainstem raphe nuclei in the human brain and is functionally similar to mouse PET1.^55,56,99^ The reported FEV mRNA is therefore most likely reflective of dorsal raphe FEV, as FEV is not expressed in any cells other than 5-HT neurons.^56^ Furthermore, congruent with findings in rodents, in patients we also observed that striatal GDNF levels are correlated with both TPH2 and SERT levels.

We observe across all three (SCZ, DP and MDD) human neuropsychiatric disorders, that striatal GDNF levels are abnormally increased in 15 of 51 patients, with significant increases in striatal GDNF observed in BP and SCZ. Of those 15 patients with abnormally elevated GDNF, 5 display a common gene expression pattern termed “GDNF/5-HT response group” which manifested independent of the condition (the group contains 3 SCZ, 1BP and 1MDD patient). The GDNF/5-HT response group may mark a mechanistic subgroup which reflects the etiological overlap suspected to exist in particular for SCZ, BP and MDD.^105–109^ As our findings suggest, both developmental as well as adult-onset abnormal GDNF upregulation associates with 5-HT system hyperstimulation and is likely to lead to or contribute to, depending on other genetic and environmental factors, specific disorder and treatment outcomes. Supporting this line of thought, increased TPH2 mRNA levels in the dorsal raphe have been observed in the brains of human patients who have committed suicide due to depression.^117^ Our results may therefore create an opportunity to devise precision medicine in the future.

Previously, we have found that abnormally high GDNF levels correlate with disease severity in first episode psychosis (FEP) patients CSF.^61^ It would be important to extend studies across the neuropsychiatric continuum and identify proxy biomarkers in the plasma to reflect CSF GDNF levels for low threshold clinical assessment. Longitudinal analysis on how the GDNF-high subgroup responds to available treatments may provide an immediate route for tailored treatment design. In parallel, repurposing inhibitors targeting RET, the main signaling receptor for GDNF in monoaminergic neurons^43–45^ currently used to treat RET positive tumors^118–120^ could be considered.

Notably a 1.5- to 2-fold increase in GDNF expression level is achieved by drugs such as methamphetamine in monkeys, rats, and mice.^47–50^ It is therefore feasible to suggest that an individual’s starting point in GDNF level and the extent of its induction may determine the susceptibility to neuropsychiatric illness; for example, individuals with low natural GDNF levels may be less prone to develop monoaminergic over-stimulation and neuropsychiatric illness upon methamphetamine abuse.

In conclusion, we show causality in mice by showing that GDNF levels regulate both 5- HT system development and adult function as well as fluoxetine response. We demonstrate increased GDNF in human neuropsychiatric disorders, and we identify a subgroup of patients with high GDNF and an aligned molecular response. The summary of our findings and proposed mechanism is depicted in Figure 8. Our experimental data suggest that a relatively small variance in GDNF expression is likely to influence normal interindividual variance in 5-HT-related behaviors. Similarly, pathological GDNF levels are likely to affect disorder symptomatology and treatment response while pharmacological modulation of GDNF signaling and monitoring its levels may offer an opportunity to modulate neuropsychiatric illness in the future.

**Figure 8.**
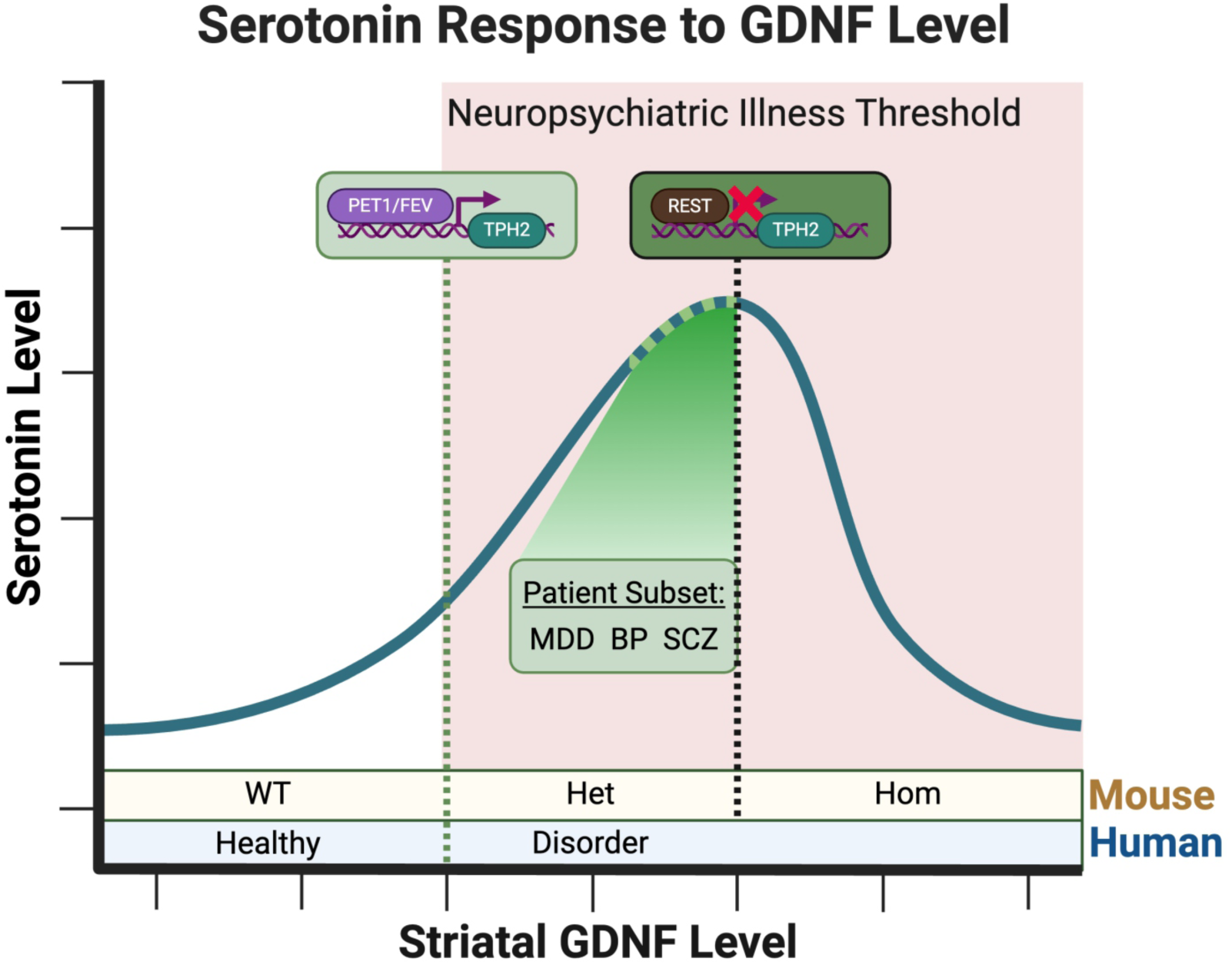
Summary Figure. As striatal GDNF increases, 5-HT levels also begin to rise with induction of 5-HT-specific transcription factor PET1/FEV. Above a certain GDNF threshold, analogous to GDNF levels observed in conditional hypermorphic heterozygous mice, only humans with neuropsychiatric illness are observed. A subset of patients displaying a common response pattern of changes to 5-HT-related genes is observable at higher levels of GDNF/5-HT induction, denoted as green dots on the 5-HT level curve. Upon further increase in GDNF level as for example in homozygous GDNF conditional hypermorphic mice, transcriptional repressor REST becomes induced, dampening 5-HT signaling resulting in an inverted U-shaped curve response to GDNF.

## 5 Resource Availability

### Lead Contact

Requests for further information and resources should be directed to and will be fulfilled by the lead contact, Jaan-Olle Andressoo

### Materials Availability

No new or unique reagents were generated from the study

### Data and code availability

The lead contact will share all data and any necessary related information required to reanalyze the data reported in this paper upon request, as well as all code used in the study.

## 6 Acknowledgements

The authors thank S. Wiss & J. Lahtinen for technical assistance, and all the members of the Andressoo lab, and especially Elina Nagaeva, for the helpful discussions and comments on the manuscript. The Helsinki Institute of Life Science and Biocenter Finland supported the Mouse Behavioral Phenotyping Facility, which the authors thank for help with behavioral assays. The authors would also like to thank BEA, the Bioinformatics and Expression Analysis core facility, which is supported by the board of research at the Karolinska Institute. JOA was supported by the Academy of Finland (grants no. 297727 and 350678), JAES Foundation grant no. 21-17628-68, Sigrid Juselius Foundation, Center of Innovative Medicine (CIMED), Hjärnfonden, Team Rynkeby-God Morgon Skolloppet grant, Åhlén-stiftelsen grant, Swedish Research Council (grants no. 2019- 01578 and 2022-01093), Helsinki Institute of Life Science, ERA-NET NEURON grant no 352077. DRG was supported by the University of Helsinki Brain and Mind doctoral program, the Fulbright Finland foundation, the Biomedicum Helsinki Foundation, and the Finnish Parkinson Foundation. VI was supported by the University of Helsinki Brain and Mind doctoral program.

## 7 Author Contributions

Conceptualization, D.R.G. and J-O.A.; Methodology, D.R.G and J-O.A.; Software, I.R., A.D., and F.F-B.; Formal Analysis D.R.G and I.R.; Investigation, D.R.G, I.R., L.C., T.S., A.R.M-R., V.I., J.J.K., and K.M.; Data Curation, I.R., A.D. and F.F-B.; Resources, T.P.P. and J-O.A.; Writing – Original Draft, D.R.G; Writing – Review & Editing, D.R.G, I.R. and J-O.A.; Visualization, D.R.G., I.R. and J-O.A.; Supervision, J-O.A.; Project Administration, J-O.A.; Funding Acquisition, J-O.A.

## 8 Declaration of Interests

The authors have no competing interests to declare.

## 9 Supplemental information

Document S1. Figures S1-S8 and Tables S1-S3

## Notes

### Competing Interest Statement

The authors have declared no competing interest.

