## Supplementary material for "Serotonin system development and adult function are regulated by GDNF": Figures S1-S8 and Tables S1-S3

**Tables S1-S3**

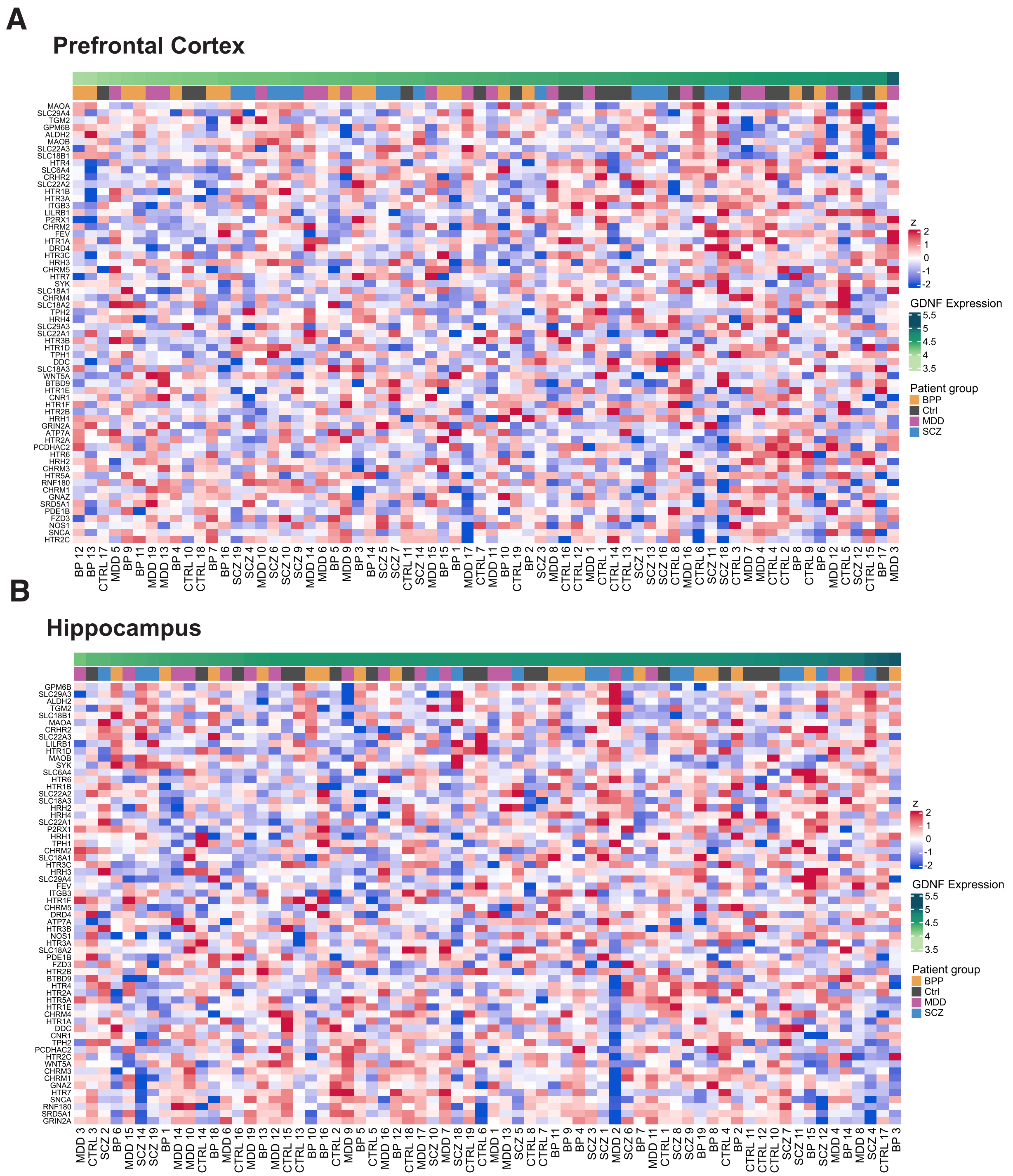


Figure S1

**Figure S1.** PFC and Hippocampal heatmaps of 5-HT gene set responses. (A) No clear patterns emerge among 5-HT gene set responses in the PFC of patients and controls from Lanz et al., 2019, with patients and controls organized by ascending levels of GDNF (B) 5-HT gene set list does not define any subsets of patients in the hippocampus with patients and controls organized by ascending levels of GDNF. In both areas, patients and controls largely fall within the same range of GDNF expression.

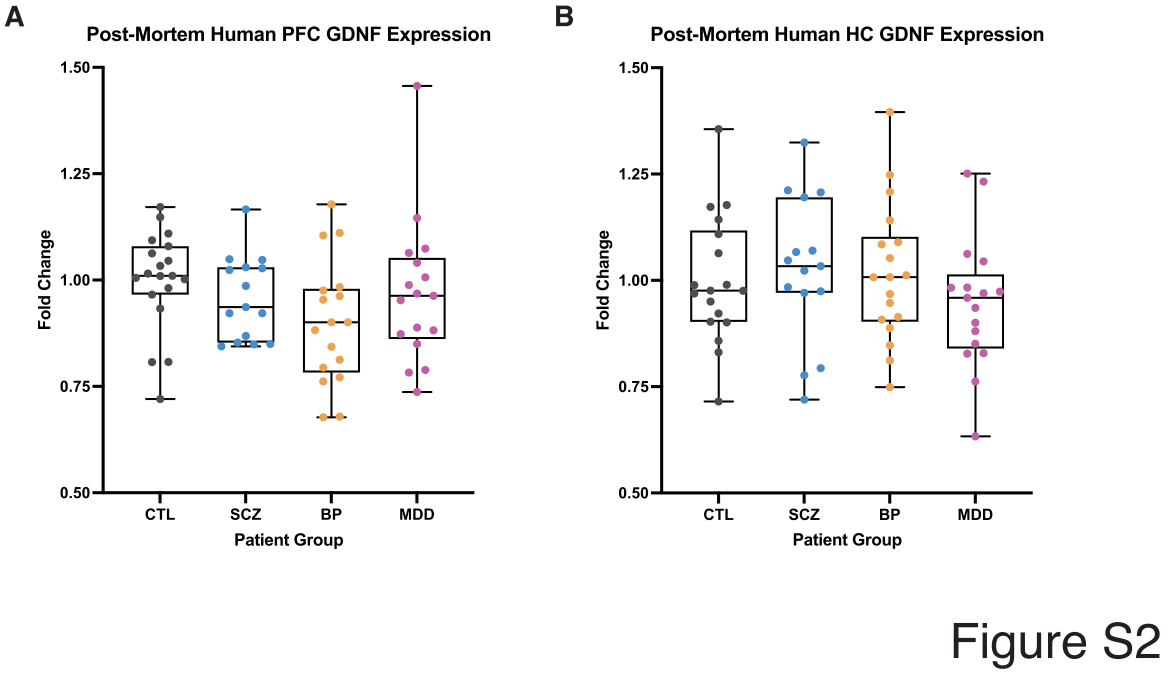


**Figure S2.** GDNF levels in the PFC and Hippocampus of human patients. (A) No significant differences in the levels of GDNF were observed in any neuropsychiatric disorder group compared to controls. (B) In the hippocampus, there were no observable differences in GDNF levels among any disorder group compared to controls.

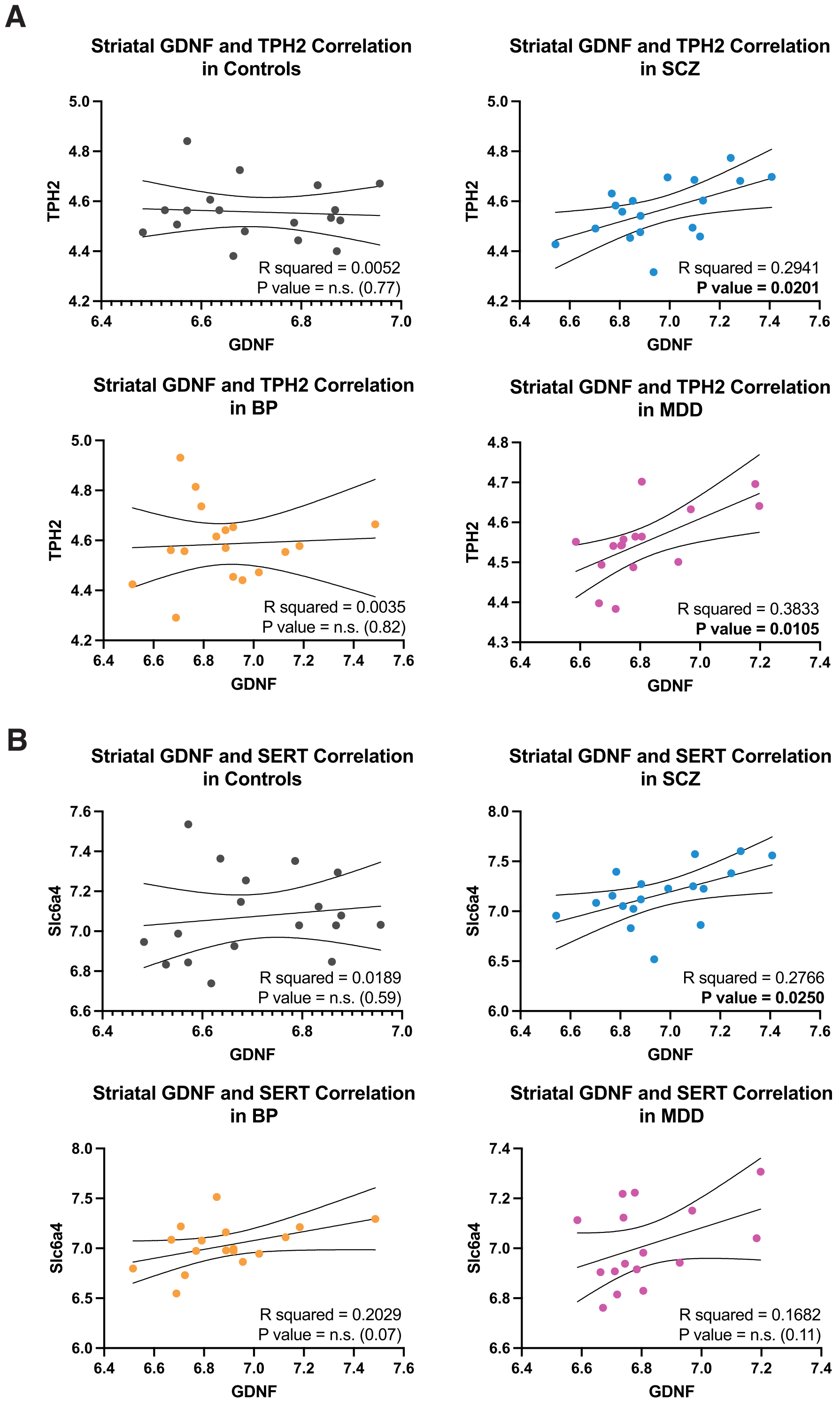


Figure S3

**Figure S3.** Subdivided correlations in each patient subgroup and controls from Lanz et al.

2019. (A) Correlations between striatal GDNF and TPH2; GDNF and TPH2 mRNA levels significantly correlate positively in MDD and SCZ patient subgroups. (B) Striatal GDNF and SERT mRNA levels also have a significant positive correlation in SCZ patients.

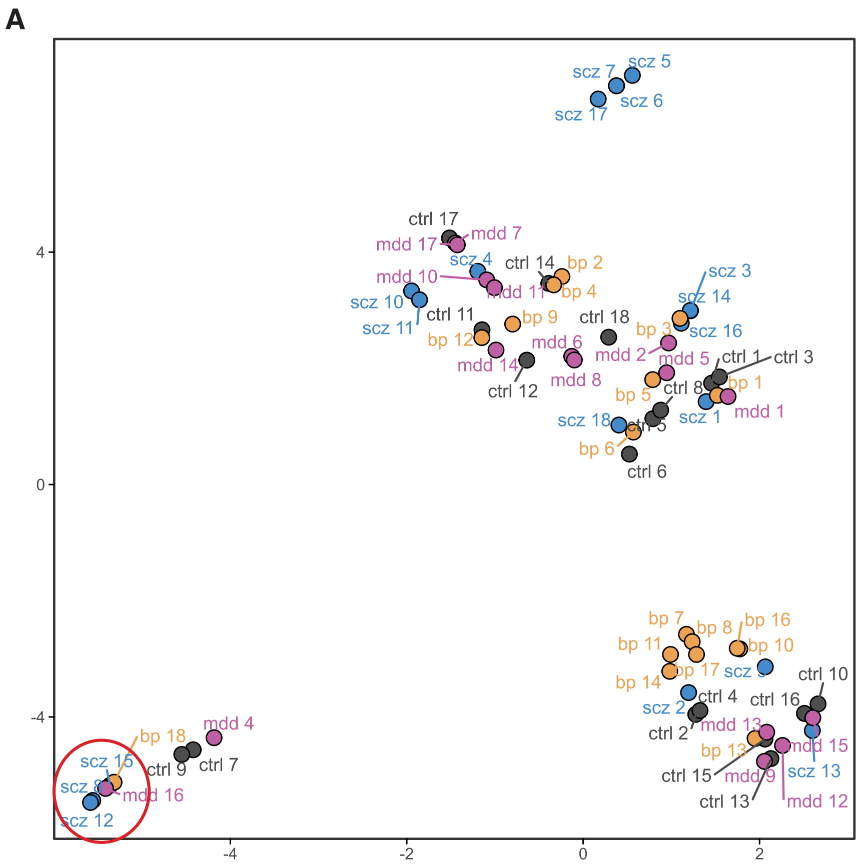


Figure S4

**Figure S4.** GDNF/5-HT response group of patients. (A) UMAP plot indicating distinct cluster of GDNF/5-HT response group patients from Lanz et al., 2019.

A

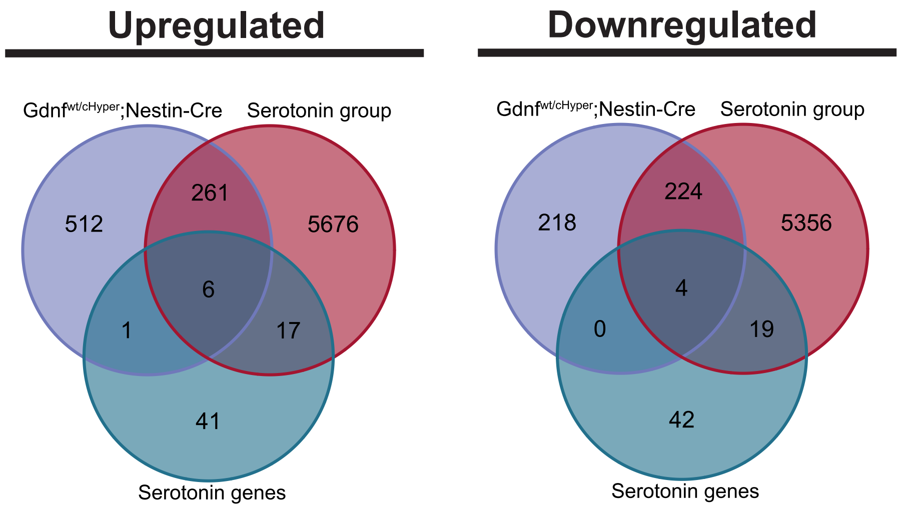


B

| Group | Overlap with 5-HT genes (FDR) |
|---|---|
| GDNF/5-HT response group STR DE genes | 3.762451e-04 |
| HET Nestin-cre STR DE genes | 5.962241e-04 |

Figure S5

**Figure S5.** Up and downregulated 5-HT genes in GDNF/5-HT response group and mutant mouse cHyper heterozygotes (A) Venn diagrams depicting significantly overlapping upregulated and downregulated 5-HT-related genes between GDNF/5-HT response group patients and Nestin-Cre GDNF wt/cHyper heterozygotes. (B) Percent of the 5-HT genes from the gene set list which are up- or downregulated in the human GDNF/5-HT response group or Nestin-Cre heterozygous cHyper mice.

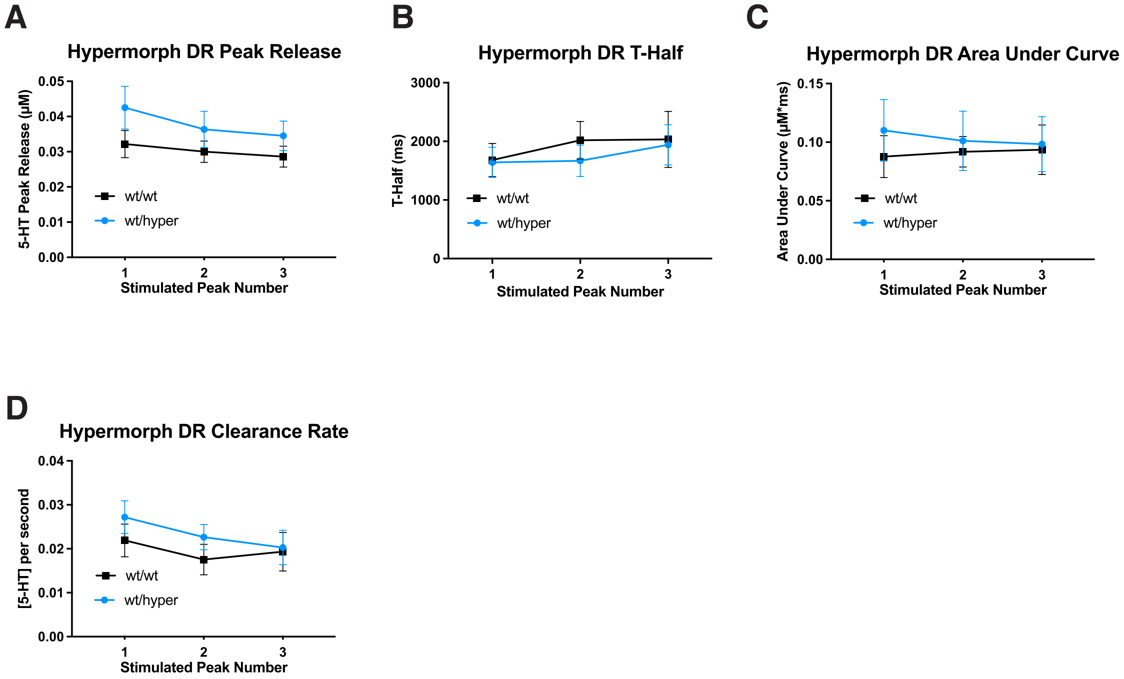


Figure S6

**Figure S6.** 5-HT release and reuptake in the dorsal raphe. (A) A nonsignificant trend towards increased stimulated 5-HT release measured by cyclic voltammetry in Dorsal Raphe slices of adult GDNF hypermorphs elicited by a 30 Hz train of electric pulses. 30-pulse, 30Hz stimulations are done every 2 minutes. (Two-way repeated measures ANOVA $p>0.05$). N=7 slices per genotype. (B) No observable difference in the time it takes for the extracellular 5-HT concentration to reach half of its peak value (t-half) in the Dorsal Raphe. (C) No observable difference in the area under the curve of the release of 5-HT elicited by a train of pulses. (D) The clearance of 5-HT in the Dorsal Raphe as measure by the peak concentration divided by the t-half was not significant across stimulations.

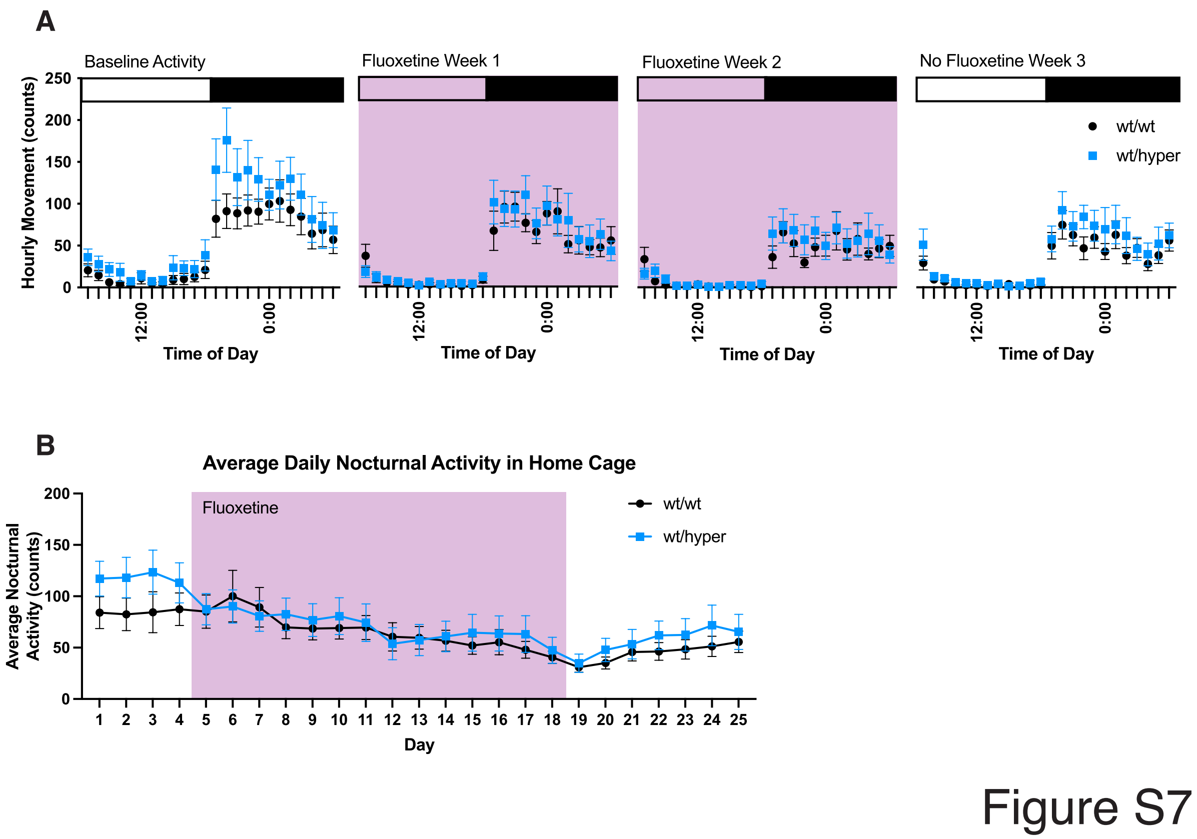


**Figure S7.** Heterozygous GDNF hypermorphic animal home cage activity. (A) Average Daily activity per hour of GDNF hypermorphic animals in home cage before and after fluoxetine. (B) Average Daily Nocturnal Activity in the Home Cage of GDNF hypermorphic and wild-type animals. 15mg/kg/day fluoxetine is given in drinking water from day 5 to day 18 (purple box). Hypermorphic animals display a nonsignificant trend towards increased baseline activity (ANOVA $p > 0.05$).

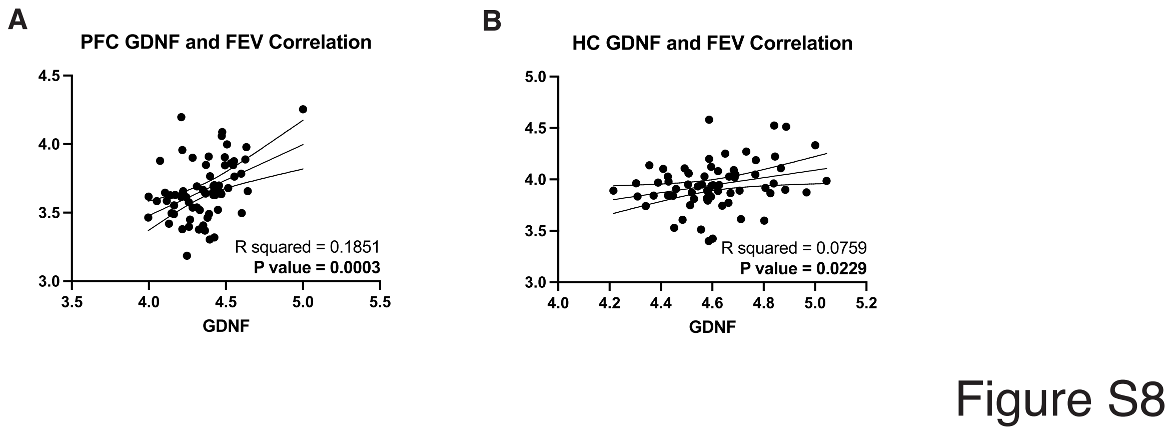


**Figure S8.** Correlations between GDNF and FEV in other brain regions from Lanz et al. 2019. In human patients and controls, GDNF and FEV display significant positive correlations in both the (A) prefrontal cortex and (B) hippocampus as assessed with the calculated Pearson correlation coefficients.

**Table S1.** A summary of studies looking at the effect of GDNF on the brain 5-HT system.

| | Paper Citation | Model Organism | GDNF Delivery | Effect on 5-HT? | PubMed ID |
|---|---|---|---|---|---|
| GDNF has a mixed effect on or increases 5-HT | Naumenko, V.S. et al., 2013 | 1. Depression-like model: ASC/Icg (Antidepressant Sensitive Cataleptics) mice<br>2. Control: CBA/Lac mice | GDNF (800 ng, i.c.v.) | Mixed; no effect in controls | PMID: 24105724 |
| | Beck, K.D. et al., 1996 | Rats, intracranial GDNF delivery | Striatal or Nigral GDNF Injections:<br>Neonatal at P8, P14, P28 (5 µg, Bilateral)<br>Adult Rats (10 µg, Unilateral) | Mixed; Neonatal Only increased 5-HT, no effect after P28 | PMID: 8785063 |
| | Ducray, A. et al., 2006 | Rat E14 ventral midbrain primary cultures | 10-100 ng/ml GDNF | Increased | PMID: 16380100 |
| | Martin, D. et al., 1996a | Adult male Fischer 344 rats | 100ng recombinant human GDNF /4uL PBS unilaterally injected intranigrally or intrastriatally | Mixed/sometimes increased, sometimes no effect | PMID: 8997607 |
| | Martin, D. et al., 1996b | Adult male Fischer 344 rats | 100ng recombinant human GDNF/4uL PBS administered i.c.v. | Increased in some areas, no effect in others | PMID: 8752595 |
| | Pertusa, M. et al., 2008 | Aged, 24-month-old male Fisher 344 rats | Intrahippocampal lentivirus injections into dorsal CA1 targetting astrocytes with GDNF transgene OE | Increased | PMID: 17399854 |
| | Galter, D., & Unsicker, K., 1999 | Primary raphe cultures from e14 rats | 5ng/ml recombinant human GDNF | Increased | PMID: 10197774 |
| GDNF has little to no effect on 5-HT | Lin, L.F. et al., 1993 | Embryonic midbrain cultures | Recobinant human GDNF overexpressed in E. coli (0.001-1000 ng/ml) | No effect | PMID: 8493557 |
| | Lin, L.F. et al., 1994 | Embryonic midbrain cultures | Purified native GDNF (0.3ng/ml) | No effect | PMID: 8035200 |
| | Hoffer, B.J., 1994 | Fisher 344 rats lesioned with 6-OHDA injection to MFB | Intranigral injection 4-weeks post-lesion (100ug) | No effect | PMID: 7891873 |
| | Gash, D. M., 1995 | Adult female rhesus monkeys | 150ug unilateral Intranigral Injection of human recombinant GDNF | No effect | PMID: 8847404 |
| | Cass W. A., 1996 | 4 injections at 2-hour intervals of methamphetamine (5mg/kg, s.c.) given to rats | 10ug intrastriatal GDNF administered 24 hhrs before methamphetamine | No effect | PMID: 8987838 |
| | Schaller, B. et al., 2005 | e18 rat ventral midbrain primary cultures | 10ng/ml GDNF added to cultures | No effect | PMID: 15725414 |
| | Hudson, J. et al., 1995 | male Fischer 344 rats | 10ug GDNF unilaterally injected to either the SN or STR and animals sacrificed 1 or 3 weeks later | Mixed/mostly no effect | PMID: 7712205 |
| | Bowenkamp, K.E. 1997 | bilaterally 6-OHDA lesioned rats | 2 i.c.v. infustions of GDNF 3 weeks apart, first 250ug, second 500ug | No effect | PMID: 9184114 |
| | Mijatovic, J. et al., 2007 | Constitutively active RET in MEN2B knock-in mice | Met918Thr mutation leading to constitutively active RET, the receptor for GDNF | No effect | PMID: 17475787 |

**Table S2.** Primers sequences used in the study for quantitative PCR analysis

| **Primers used for qPCR** | | |
|---|---|---|
| **Primer** | **Forward Sequence** | **Reverse Sequence** |
| Gdnf | CGCTGACCAGTGACTCCAATATGC | TGCCGCTTGTTTATCTGGTGACC |
| Tph2 | AGAGTTGGAGACGGAGTCGT | AAGGGCAGTGGCTTATGACC |
| Slc6a4 (Sert) | CCCAGACTCTTGTGGGTTCC | CTAGCTGATGACTGGGTGGC |
| Pet1/Fev | CATGTACCTGCCAGATCCCG | GGAGAAACTGCCACAACTGG |
| Rest | GCACAGTTCAGAGGAGTACAG | CCCATGTTGGCACTGTTGTT |
| Actin | CTAAGGCCAACCCTGAAAAG | ACCAGAGGCATACAGGGACA |
| Gapdh | GCCTCGTCCCGTAGACAAAA | ATGAAGGGGTCGTTGATGGC |
| Rn18s | CTTAGAGGGACAAGTGGCG | ACGCTGAGCCAGTCAGTGTA |

**Table S3.** Primer sequences used in the study for genotyping

| **Primers used for Genotyping** | |
|---|---|
| **Primer** | **Sequence** |
| GDNFh wt F | GAA ACC AAG GAG GAA CTG ATC |
| GDNFh mut F1 | CGG TGG GCT CTA TGG CTT CT |
| GDNFh R | TCT TCT GCC TCT GCC TCC G |
| cHyper F | TCT AAG AAA GCA TTC CGC TAA ACG |
| cHyper R1-1 | TTC CAG GGT CAA GGA AGG CAC |
| cHyper R2-1 | GGA TGC GGT GGG CTC TAT G |
| cHyper R3-1 | TCC GCC ATC TTG GTC CTT ATC |
| Cre 3’ | CGT TTT CTG AGC ATA CCT GGA |
| Cre 5’ | AAT CTC CCA CCG TCA GTA CG |
| GDNF 5’ wt F | CTC ATT TCC CAC AGG GAA CTG |
| GDNF 3’ wt F | GAA ACC AAG GAG GAA CTG ATC |
| GDNF 3’ wt R | TCT TCT GCC TCT GCC TCC G |
